# ARID1A maintains transcriptionally repressive H3.3 associated with CHD4-ZMYND8 chromatin interactions

**DOI:** 10.1101/2022.02.27.482165

**Authors:** Jake J. Reske, Mike R. Wilson, Brooke Armistead, Cristina Perez, Joel Hrit, Marie Adams, Scott B. Rothbart, Stacey A. Missmer, Asgerally T. Fazleabas, Ronald L. Chandler

## Abstract

ARID1A is a signature subunit of the mammalian SWI/SNF (BAF) chromatin remodeling complex and is mutated at a high rate in malignancies and benign diseases originating from the uterine endometrium. Through genome-wide analysis of human endometriotic epithelial cells, we show that more than half of ARID1A binding sites are marked by the variant histone H3.3, including active regulatory elements. ARID1A loss leads to H3.3 depletion at ARID1A bound active regulatory elements and a concomitant redistribution of H3.3 towards genic elements. ARID1A interactions with the repressive chromatin remodeler CHD4 (NuRD) are associated with H3.3-containing chromatin regulation. ZMYND8, the CHD4-interacting acetyl-histone H4 reader, specifies ARID1A-CHD4-H3.3 target regulatory activity towards histone H4 lysine 16 acetylation (H4K16ac) to repress super-enhancers. ARID1A, H3.3, CHD4, and ZMYND8 co-repress the expression of genes governing extracellular matrix, motility, adhesion, and epithelial-to-mesenchymal transition. Moreover, these gene expression alterations are observed in human endometriomas. Altogether, these studies demonstrate that cooperation among a histone reader and different types of chromatin remodelers safeguards the endometrium through transcriptionally repressive H3.3.

## Introduction

The SWI/SNF complex remodels chromatin through ATP-dependent DNA sliding, H2A/H2B dimer eviction, and nucleosome ejection functions (Dechassa et al., 2010; Kassabov, Zhang, Persinger, & Bartholomew, 2003; Whitehouse et al., 1999). SWI/SNF remodeling activities open chromatin and promote accessibility for other DNA-binding factors and chromatin regulators (Clapier, 2021; Clapier & Cairns, 2009). SWI/SNF complex composition is heterogeneous and cell type dependent (Wang et al., 1996). SWI/SNF regulates lineage-specific enhancer activity through multiple mechanisms (Alver et al., 2017). Protein subunit architecture contributes to SWI/SNF complex specificity through specialized cofactor interactions. The activities of chromatin remodelers and associated machinery are known to modulate the epigenome by regulating histone post-translational modifications and nucleosome composition (Clapier, 2021). Multiple chromatin remodeler complexes are often observed at the same genomic loci and can perform redundant, cooperative, or antagonistic transcriptional regulatory roles (Morris et al., 2014).

Subunits within the mammalian SWI/SNF (BAF) chromatin remodeler complex are mutated across an estimated 20% of all human cancers (Kadoch et al., 2013). Tissue-specific propensities for mutations in certain SWI/SNF subunits are also evident (Kadoch & Crabtree, 2015). ARID1A (BAF250A) is the most frequently mutated SWI/SNF subunit (Mittal & Roberts, 2020). ARID1A is the largest SWI/SNF subunit and acts as a structural scaffold for other subunits in certain SWI/SNF complexes (He et al., 2020; Mashtalir et al., 2018). ARID1A also exhibits essential DNA-binding activity albeit in a non-sequence-specific manner (Chandler et al., 2013; Dallas et al., 2000). Defects in chromatin accessibility and higher-order chromatin structure are thought to underlie ARID1A and SWI/SNF mutant pathogenesis at least partially (Barutcu et al., 2016; Kelso et al., 2017). Uterine endometrial cancer displays high rates of ARID1A mutation, with roughly 40% of cases showing loss of ARID1A expression (Cancer Genome Atlas Research et al., 2013; Wu, Wang, & Shih Ie, 2014). ARID1A mutations and loss of expression are also observed in deeply invasive forms of endometriosis, which is characterized by ectopic spread of the endometrium (Anglesio et al., 2017; Samartzis et al., 2012; Zondervan, Becker, & Missmer, 2020). ARID1A mutations are also common in endometriosis-associated ovarian cancers (Jones et al., 2010; Wiegand et al., 2010).

In the endometrial epithelium, we have previously shown that ARID1A normally promotes epithelial identity by repressing the expression of mesenchymal and invasion genes, at least in part through promoter-proximal and distal chromatin interactions that affect transcriptional activity (Reske et al., 2020; Wilson et al., 2020; Wilson et al., 2019). Other reports have demonstrated that ARID1A and SWI/SNF can function as a repressor, often through interactions with repressive machinery (Bui et al., 2019; Chandler et al., 2013; Rafati et al., 2011; Van Rechem, Boulay, & Leprince, 2009). Although nucleosome structure and histone post-translational modifications are suspected mechanisms, it remains poorly understood how SWI/SNF governs the epigenome.

Here, we reveal a mechanism by which ARID1A maintains histone variant H3.3 in active chromatin that is associated with biochemical and genomic interactions with the SWI/SNF-like CHD4 (NuRD) remodeler complex. This regulation is further specified toward repression of transcriptional hyperactivation through CHD4 interactions with the multivalent histone reader ZMYND8 to target H4K16 acetylated regions of the genome, including a subset of super-enhancers. We finally reveal that this mechanism of ARID1A, H3.3, CHD4, and ZMYND8 co-repression targets pathophysiological genes involved in epithelial-to-mesenchymal transition (EMT) and cellular invasion, and these genes are aberrantly upregulated in human endometriomas. Altogether, our studies reveal a role for ARID1A-containing SWI/SNF complexes in the maintenance of H3.3, and, at a subset of physiologically relevant target genes, ARID1A-CHD4-ZMYND8 interactions govern transcriptionally repressive H3.3.

## Results

### ARID1A mutations in cancer are associated with reduced histone H3.3

To reveal potential molecular mechanisms governing chromatin regulation by ARID1A in the endometrium, we leveraged the Cancer Cell Line Encyclopedia (CCLE) global chromatin profiling data set of bulk histone H3 peptide measurements by mass spectrometry (Ghandi et al., 2019). Among the 896 total cancer cell lines that were assayed along with genetic mutation profiling, 27 are specifically endometrial cancer. Cancer cell lines were segregated by ARID1A mutation status, and relative bulk H3 peptide abundances were compared between ARID1A mutant and wild-type cancer cell lines (Figure 1A). This resource and analysis allowed us to identify histone peptide abundances associated with ARID1A mutation. Across cancer and specifically within endometrial cancer, we observed that ARID1A mutant lines showed overall higher levels of H3K27ac1K36me0 and H3K79me2 and lower levels of H3K4me1, H3K4me2, and H3.3 (H3.3K27me0K36me0) (Figure 1B). Consistent with prior literature, SWI/SNF (BAF) was shown to biophysically and functionally regulate H3K4me1-marked chromatin (Local et al., 2018). Histone H3.3 is a variant of canonical H3 with known ties to active chromatin and transcriptional regulation (Ahmad & Henikoff, 2002; Shi, Wen, & Shi, 2017; Szenker, Ray-Gallet, & Almouzni, 2011). Like SWI/SNF, H3.3 has also been observed to mark and regulate active enhancers (Chen et al., 2013; Deaton et al., 2016; Martire et al., 2019; H. Zhang et al., 2017). In contrast with H3.3, abundance of the respective canonical H3 (H3.1) peptide was not different between ARID1A mutant and wild-type lines (Figure 1C-D). To rule out the possibility that cancer-associated histone gene mutations could cause this phenomenon, we recapitulated these H3.3 associations in cell lines that are wild-type for all 74 human histone genes (Nacev et al., 2019) (Figure 1—figure supplement 1). While a minor effect size, the association between ARID1A mutation and loss of H3.3 suggests that H3.3 chromatin deposition or stabilization might be mediated by ARID1A-SWI/SNF remodeler activity, and H3.3 disruption may be linked to pathogenic chromatin alterations following ARID1A mutation.

**Figure 1.**
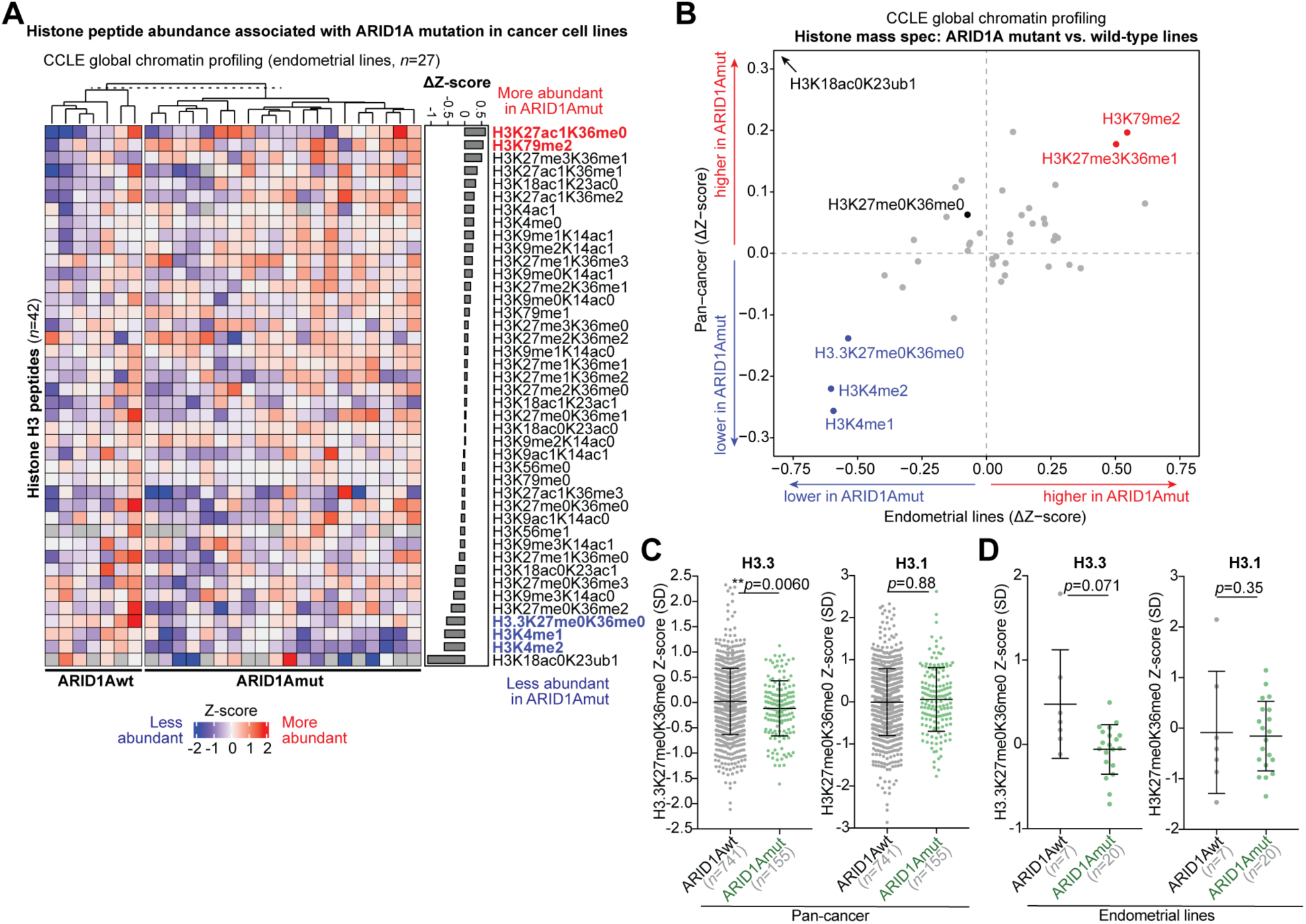
Histone peptide abundance associated with ARID1A mutation in cancer cell lines. **A**, Representation of 42 distinct histone H3 peptide measurements by mass spectrometry in public CCLE global chromatin profiling data, specifically for endometrial cancer cell lines stratified by ARID1A mutation status. Heatmap values are relative peptide abundance Z-scores, where 0 (white) is the mean across all pan-cancer cell lines. H3 peptides were then ranked (*y*-axis) by differential abundance between ARID1A mutant vs. wild-type lines. **B**, Summary of ARID1A mutant peptide associations across all cancer cell lines (*n* = 896, *y*-axis) or specifically endometrial cancer lines (*n* = 27, *x*-axis). **C**-**D**, Box dot plots showing relative abundance of H3.3 vs. canonical H3.1 peptides in ARID1A wild-type and mutant lines: **C**, pan-cancer lines; **D**, endometrial lines. Statistic is two-tailed, unpaired Welch’s *t*-test.

### ARID1A regulates H3.3-associated active chromatin

We next interrogated the chromatin regulatory roles of H3.3 in immortalized 12Z human endometriotic epithelial cells. Our previous studies have demonstrated that ARID1A promotes epithelial characteristics in 12Z endometriotic epithelial cells at both transcriptional and phenotypic levels, such that ARID1A loss leads to epithelial-to-mesenchymal transition (EMT) and enhanced migration and invasion (Wilson et al., 2020; Wilson et al., 2019). ARID1A loss in 12Z recapitulates many of the molecular and cellular features observed in ARID1A-deficient endometrial epithelia *in vivo* (Wilson et al., 2020; Wilson et al., 2019). Therefore, 12Z cells represent a model system to explore physiological roles for ARID1A in epigenomic regulation.

To measure genome-wide H3.3 localization, we performed H3.3 chromatin immunoprecipitation followed by sequencing (ChIP-seq) in control 12Z cells (*n* = 2 IP replicates). Significant H3.3 enrichment was observed at 40,006 genomic regions (Figure 2A). Intronic, intergenic, and promoter regions comprised the vast majority of H3.3 enrichment sites (Figure 2A), which may be expected as H3.3 is known to mark active regulatory elements such as enhancers and gene promoters (Chen et al., 2013). H3.3 ChIP-seq peaks were 1830 bp in width on average and ranged from <500 bp to >10 kilobases (Figure 2B). Intersecting H3.3 ChIP-seq peaks with our previously published ARID1A ChIP-seq data from these cells (Wilson et al., 2019) revealed that over half of each peak set overlapped, suggesting widespread co-regulation (Figure 2C).

**Figure 2.**
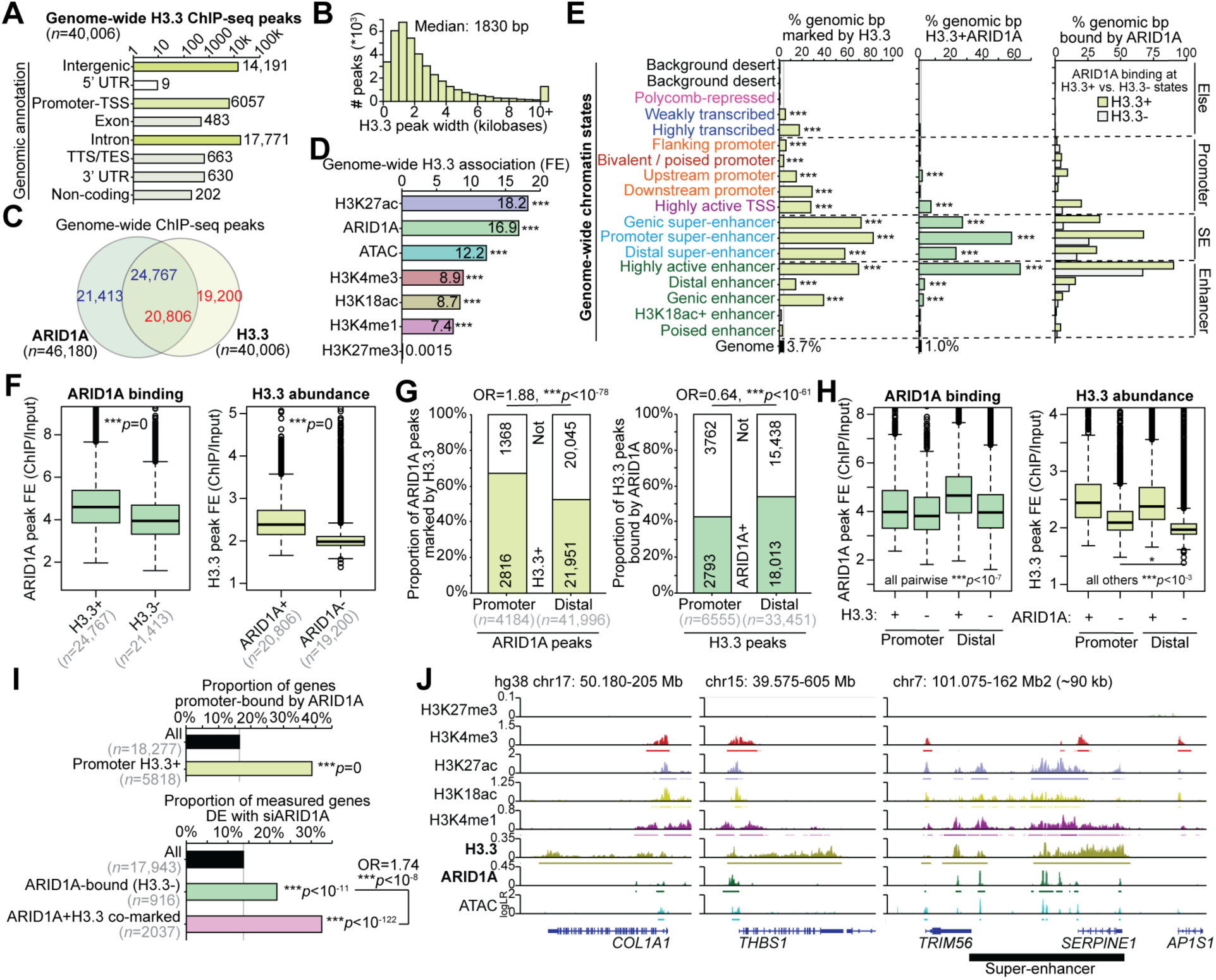
Genome-wide analysis of H3.3-ARID1A chromatin co-regulation. **A**, Genomic annotation of 40,006 genome-wide H3.3 ChIP-seq peaks in 12Z cells. **B**, Distribution of H3.3 peak widths. Median H3.3 peak width is 1830 bp. **C**, Genome-wide overlap of ARID1A and H3.3 ChIP-seq peaks. **D**, Genome-wide association between H3.3 and other previously measured chromatin features, per genomic bp. Statistic is hypergeometric enrichment. **E**, Enrichment for H3.3 and ARID1A co-regulation across 18 chromatin states previously modeled via *ChromHMM* (Wilson et al., 2020). Left, enrichment of H3.3 peaks; center, enrichment of H3.3+ARID1A binding; right, enrichment of ARID1A binding at sites with vs. without H3.3. Statistic is hypergeometric enrichment. **F**, Left, ARID1 A binding levels (ChIP/input fold-enrichment, FE) at H3.3+ vs. H3.3-ARID1A peaks. Right, H3.3 abundance (ChIP/input fold-enrichment) at ARID1A+ vs. ARID1A-H3.3 peaks. Statistic is two-tailed, unpaired Wilcoxon’s test. **G**, Association between ARID1A and H3.3 co-binding at promoter proximal (<3 kb from a TSS) vs. distal (>3 kb from a TSS) peaks. Statistic is two-tailed Fisher’s exact test. **H**, Left, ARID1A binding levels at promoter vs. distal peaks with or without H3.3. Right, H3.3 abundance at promoter vs. distal peaks with or without ARID1A. Statistic is two-tailed, unpaired Wilcoxon’s test. **I**, Top, enrichment of H3.3 at genes promoter-bound by ARID1A. Bottom, enrichment of ARID1A+H3.3 co-binding at genes DE following ARID1A loss (siARID1A treatment). Statistics are hypergeometric enrichment test and pairwise two-tailed Fisher’s exact test. **J**, Example hg38 browser shots of genes and regulatory elements co-regulated by H3.3 and ARID1A. *y*-axis is log-likelihood ratio (logLR) of assay signal (compared to input chromatin for ChIP-seq or background genome for ATAC-seq). Small bars under tracks indicate significant peak detection by *MACS2* (FDR < 0.05). Super-enhancers were detected by *ROSE* from H3K27ac ChIP-seq. * *p* < 0.05, ** *p* < 0.01, *** *p* < 0.001.

We previously constructed a genome-wide chromatin state map accompanying ARID1A loss in 12Z cells via *chromHMM* (Ernst & Kellis, 2012) by measuring seven chromatin features associated with transcriptional regulation: total RNA, ATAC (accessibility), H3K27ac, H3K18ac, H3K4me1, H3K4me3, and H3K27me3 (Wilson et al., 2020). Similar to our previous reports of ARID1A regulated chromatin states, genomic H3.3 enrichment was highly associated with all active, euchromatic features, but not heterochromatic H3K27me3 (Figure 2D). Annotating H3.3 enrichment in each of our characterized chromatin states revealed that H3.3 is associated with similar regulatory chromatin states as ARID1A binding, most notably super-enhancers and active typical enhancers (Figure 2E, left). In agreement, co-regulation by H3.3 and ARID1A was most prominently observed at these same chromatin states (Figure 2E, center). Next, we examined ARID1A binding at H3.3-marked vs. H3.3-absent chromatin sub-states and found that ARID1A binding was associated with H3.3 at promoter and genic super-enhancers and active transcription start sites (TSS) (Figure 2E, right).

Upon further investigation of ARID1A and H3.3 genome-wide binding patterns, we observed that genome-wide ARID1A peaks showed overall stronger ARID1A binding when H3.3 was also localized, and H3.3 was overall more abundant at genome-wide H3.3 peaks also bound by ARID1A (Figure 2F). ARID1A and H3.3 ChIP-seq peaks were then broadly classified into promoter-proximal (located within 3 kb of a TSS) and distal (located further than 3 kb from a TSS). Interestingly, promoter-proximal ARID1A peaks were more likely to show H3.3 co-marking than distal ARID1A peaks, whereas distal H3.3 peaks were more likely to show ARID1A co-binding than promoter-proximal H3.3 peaks (Figure 2G). Further, ARID1A and H3.3 peaks were segregated into four classes based on binding status at promoter vs. distal sites. ARID1A binding was strongest at distal peaks marked by H3.3, while H3.3 was most abundant at promoter-proximal peaks bound by ARID1A (Figure 2H). These genome-wide binding patterns indicate that ARID1A and H3.3 may co-regulate active chromatin elements like enhancers and promoters.

We previously reported that ARID1A chromatin binding near gene promoters is associated with transcriptional regulation, such that ARID1A loss leads to aberrant gene expression (Wilson et al., 2019). Our H3.3 data further revealed that ARID1A promoter binding is highly enriched among genes marked by promoter H3.3 (Figure 2I, top), indicating that ARID1A transcriptional regulation may be coupled with H3.3. Moreover, the 2037 genes co-marked by ARID1A and H3.3 in the promoter region were more likely to show differential expression (DE) following ARID1A loss than genes without promoter H3.3 (Figure 2I, bottom). In addition, locus-scale investigation clearly showed that ARID1A and H3.3 often co-mark active chromatin regulatory elements, which infrequently also includes gene body coating by H3.3, such as at *COL1A1*, *THBS1*, and *SERPINE1* (Figure 2J). These data collectively suggest H3.3 may be linked to transcriptional regulatory activity by ARID1A at the level of chromatin.

### ARID1A chromatin interactions maintain H3.3

To understand the relationship between ARID1A and H3.3, we depleted ARID1A from 12Z cells using lentiviral shRNA particles targeting ARID1A (shARID1A) then measured H3.3 by ChIP-seq. Our differential H3.3 ChIP-seq analysis (shARID1A vs. non-targeting shRNA control, *n* = 2) indicated that nearly 1/3 of tested H3.3 regions showed significant differences in H3.3 abundance (*csaw*/*edgeR*, FDR < 0.05) at 72 hours following ARID1A knockdown (Figure 3A, Figure 3—figure supplement 1A). We noted that ARID1A knockdown in 12Z cells did not result in obvious changes in global H3.3 levels by immunoblotting (Figure 3—figure supplement 1B), suggesting any effects are likely occurring at the level of chromatin. This is further supported by our previously reported 12Z ARID1A knockdown RNA-seq data (Wilson et al., 2019) indicating that the dominantly expressed H3.3-encoding gene isoform, *H3F3B*, does not change in expression (Figure 3—figure supplement 1C). Genomic regions that showed significantly affected H3.3 following ARID1A loss had overall lower baseline levels of H3.3 compared to stable H3.3 regions that were not affected by ARID1A loss (Figure 3B). We then investigated how ARID1A chromatin binding may be directly associated with the observed changes in H3.3 following ARID1A loss. Strikingly, ARID1A-bound differential H3.3 regions almost exclusively lost H3.3 and rarely gained H3.3 (Figure 3C-D). While 33% of all tested H3.3 regions had detectable ARID1A binding, 81% of the 8418 shARID1A decreasing H3.3 regions were normally bound by ARID1A, as opposed to only 3% of the 11,059 shARID1A increasing H3.3 regions (Figure 3D). These results indicate that ARID1A interactions with H3.3 chromatin may serve to promote H3.3 incorporation or maintain its stability.

**Figure 3.**
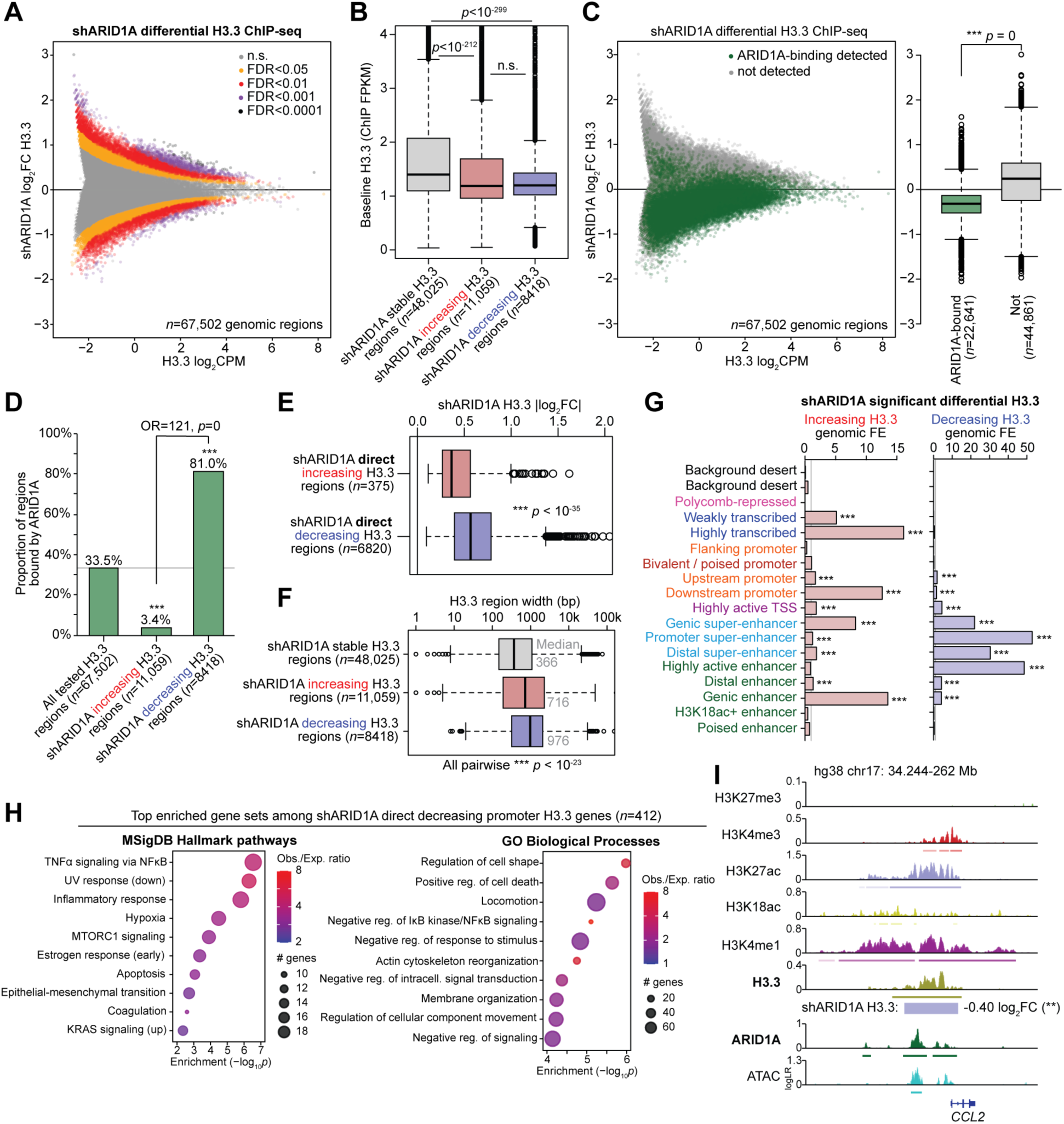
Genome-wide analysis of ARID1A-dependent H3.3. **A**, MA plot of shARID1A vs. control differential H3.3 ChIP-seq, *n* = 67,502 tested genomic regions. Regions are colored based on shARID1A differential H3.3 significance. FDR < 0.05 was used as the significance threshold for all downstream analyses. **B**, Baseline H3.3 abundance (ChIP FPKM) at regions with stable H3.3 following ARID1A knockdown, regions that display increasing H3.3, and regions that display decreasing H3.3. Statistic is two-tailed, unpaired Wilcoxon’s test. **C**, shARID1A differential H3.3 regions segregated by detection of ARID1A binding in wild-type cells. Left, MA plot with all genome-wide H3.3 tested regions, colored by ARID1A binding status. Right, box plot quantification of shARID1A log_2_FC H3.3 abundance, segregated by ARID1A binding status. Statistic is two-tailed, unpaired Wilcoxon’s test. **D**, Enrichment of ARID1A binding detection at regions with decreasing H3.3 following ARID1A loss compared to all tested H3.3 regions. Statistics are hypergeometric enrichment test and pairwise two-tailed Fisher’s exact test. **E**, Magnitude of H3.3 change (log_2_FC) among ARID1A-bound (direct) shARID1A significantly decreasing vs. increasing H3.3 regions. Statistic is two-tailed, unpaired Wilcoxon’s test. **F**, Distribution of H3.3-enriched region widths among shARID1A stable vs. increasing vs. decreasing H3.3 regions. Statistic is two-tailed, unpaired Wilcoxon’s test. **G**, Chromatin state enrichment among shARID1A increasing and decreasing H3.3 regions, calculated per 200 bp genomic interval. Statistic is hypergeometric enrichment. **H**, Top 10 significant (FDR < 0.05) enriched Hallmark pathways (left) and GO Biological Process gene sets (right) among genes with shARID1A direct (ARID1A-bound) decreasing promoter H3.3. **I**, Representative hg38 locus near *CCL2* displaying H3.3 maintained by ARID1A chromatin interactions. *** *p* < 0.001.

We further characterized the changes in H3.3 occurring following ARID1A loss. Globally, we found that typical enhancers (distal regions marked by H3K27ac and ATAC, >3 kb away from a TSS and excluding super-enhancers) were enriched for shARID1A-driven H3.3 alterations as compared to gene promoters and super-enhancers (Figure 3—figure supplement 1D). Intriguingly, gene promoters displayed both decreasing and increasing H3.3, whereas typical enhancers and super-enhancers almost exclusively lost H3.3 if significantly affected (Figure 3—figure supplement 1E). Among genomic H3.3 regions that were directly affected by ARID1A loss, decreasing H3.3 regions tended to display greater differences in H3.3 abundance than increasing H3.3 regions (Figure 3E). Regions that displayed decreasing H3.3 also tended to have overall wider genomic footprints than increasing or stable H3.3 regions (Figure 3F). In agreement with where ARID1A-H3.3 co-regulation is most frequently observed, chromatin state enrichment analysis indicated that ARID1A loss led to depletion of H3.3 at promoter super-enhancers and highly active enhancers, while increasing H3.3 was observed over actively transcribed gene bodies (Figure 3G). From the 412 genes we identified with shARID1A direct decreasing promoter H3.3, we found significant enrichment for inflammatory, hypoxia, apoptosis, locomotion, and EMT pathways, such as *CCL2* (Figure 3H-I). These data suggest that ARID1A maintains H3.3 at active regulatory elements such as enhancers and super-enhancers, and, when ARID1A is lost, redistribution of H3.3 occurs towards active genes already marked by H3.3.

### H3.3 depletion phenocopies transcriptional effects of ARID1A loss

We next sought to determine the transcriptional consequences of H3.3 loss in endometrial epithelia. We hypothesized that *H3F3B* could be knocked down to reduce H3.3 levels for acute transcriptome evaluation without impeding cell health (Figure 4A). Using siRNA targeting *H3F3B* (siH3F3B), we observed H3.3 depletion by immunoblotting without affecting the cell cycle (Figure 4B, Figure 4—figure supplement 1A-B). RNA-seq transcriptome analysis 72 hours following siRNA transfection showed clear loss of *H3F3B* expression, but not *H3F3A*, accompanying 1608 significant DE genes (*DESeq2*, FDR < 0.001) including those both upregulated (repressed by H3.3) and downregulated (activated by H3.3) (Figure 4C-E). As expected, we also observed highly significant enrichment for H3.3 dependent transcriptional changes among genes marked by promoter H3.3 (Figure 4—figure supplement 1C). Similar to our previous observations with acute ARID1A loss (Wilson et al., 2019), depletion of H3.3 led to mostly minor alterations in gene expression, with the majority of DE genes displaying <0.5 log_2_FC expression change (Figure 4E). These data indicate H3.3 serves both activating and repressing roles in transcriptional regulation of endometrial epithelial cells.

**Figure 4.**
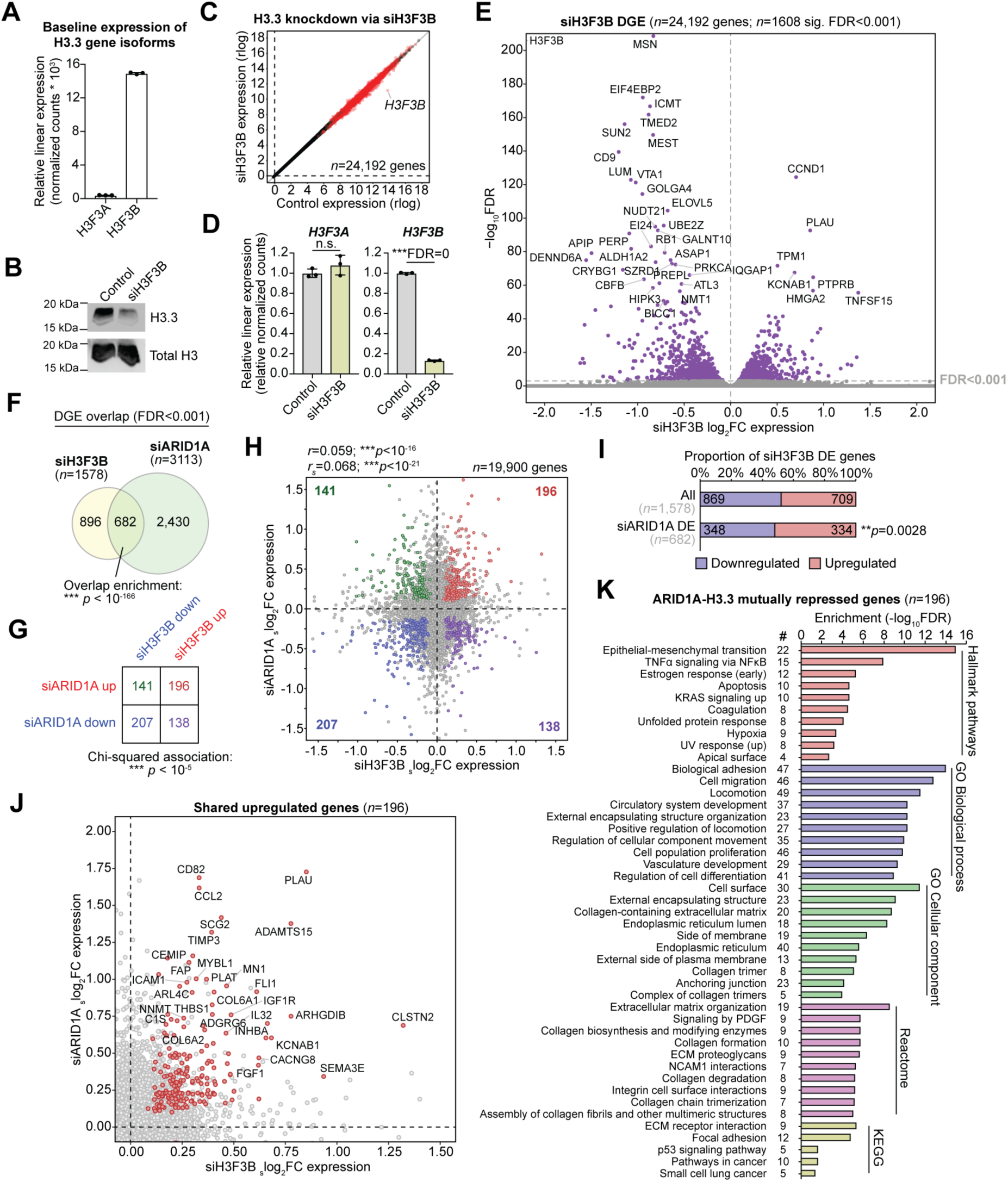
Transcriptional effects of H3.3 depletion and overlap with ARID1A. **A**, Baseline relative linear expression of *H3F3A* (*H3-3A*) and *H3F3B* (*H3-3B*) gene isoforms encoding H3.3, as measured by RNA-seq. **B**, Western blot for H3.3 and total H3 in control vs. siH3F3B treated cells. **C**, Global transcriptomic effects of 24,192 genes following H3.3 knockdown via siH3F3B treatment. Red dots represent significant DE genes (*DESeq2*, FDR < 0.001). **D**, Relative linear expression of *H3F3A* and *H3F3B* by RNA-seq in control and siH3F3B cells. **E**, Volcano plot depicting siH3F3B vs. control differential gene expression (DGE). Top significant genes are labeled. **F**, Significant overlap in DE genes following H3.3 knockdown (siH3F3B) vs. ARID1A knockdown (siARID1A). Statistic is hypergeometric enrichment. **G**, Directional segregation of siH3F3B/siARID1A overlapping DE genes. A positive association is observed by Chi-squared test, i.e., genes are more likely to be upregulated or downregulated in both conditions as opposed to antagonistic regulation. **H**, Scatter plot of siH3F3B vs. siARID1A expression log_2_FC (with shrinkage correction) for all 19,900 transcriptome-wide commonly detected genes. Statistic is pearson (*r*) and Spearman (*r*_s_) correlation coefficients. Colored dots indicate significant DE genes (FDR < 0.001) in both treatment conditions. **I**, Enrichment for H3.3 repression (siH3F3B upregulation) among siH3F3B genes which are also affected by ARID1A loss (siARID1A). Statistic is hypergeometric enrichment test. **J**, Scatter plot of 196 shared DE genes upregulated following knockdown of either H3.3 and ARID1A. These genes are mutually repressed by H3.3 and ARID1A. **K**, Top significant (FDR < 0.05) enriched gene sets among the 196 ARID1A-H3.3 mutually repressed genes among various gene set databases.

Comparing the gene expression changes following H3.3 loss with those following ARID1A loss, we observed significant overlap, with 682 shared dysregulated genes (Figure 4F). These 682 genes were then grouped by direction of change (upregulated vs. downregulated) to identify genes with the same or different expression patterns following ARID1A vs. H3.3 loss. A significant association was observed between the effects of H3.3 and ARID1A loss indicating shared transcriptional consequences (Figure 4G). Gene expression changes also positively correlated transcriptome-wide (Figure 4H). Intriguingly, the 682 genes affected by loss of H3.3 and ARID1A were more likely to be repressed by H3.3 (Figure 4I). 196 genes were identified as mutually repressed by both ARID1A and H3.3, including *PLAU*, *ADAMTS15*, *C1S*, *CD82*, *CCL2*, and *CLSTN2* (Figure 4J). In agreement with differential H3.3 patterns, these 196 co-repressed genes were enriched for similar gene sets as observed among the shARID1A direct decreasing promoter H3.3 gene set, including EMT, TNFα signaling, estrogen response, apoptosis, adhesion, migration, extracellular matrix, and collagens (Figure 4K). Altogether, these data suggest that ARID1A and H3.3 co-regulate similar target genes in endometrial epithelial cells. At the chromatin level, depletion or destabilization of H3.3 as a result of ARID1A loss may lead to the upregulation of a physiologically relevant set of EMT and invasion genes.

### ARID1A interacts with CHD4 to co-regulate H3.3 with ZMYND8

While our data implicate H3.3 in the ARID1A mutant endometrium, few reports have linked SWI/SNF activity to H3.3 containing nucleosomes (Gehre et al., 2020; Pillidge & Bray, 2019). To gain insight into possible chromatin regulators mediating H3.3 regulation by ARID1A, we used the ReMap2020 database of 165 million peak regions extracted from genome-wide binding assays (Cheneby et al., 2020). For all 1135 transcriptional regulators included in this database, we calculated genome-wide associations for each set of factor peaks with H3.3-marked (H3.3+) vs. H3.3-absent (H3.3-) ARID1A binding. This analysis revealed that two zinc finger MYND-type proteins, ZMYND11 (BS69) and ZMYND8 (PRKCBP1, RACK7), were among the top co-regulators associated with H3.3+ ARID1A chromatin binding (Figure 5A), suggesting that H3.3 regulation by ARID1A may be mediated by these co-regulators. ZMYND11 and ZMYND8 are multivalent chromatin readers that are suggested to function as interfaces between histones and other chromatin regulator complexes like remodelers, writers, and erasers (Guo et al., 2014; Savitsky et al., 2016). Both proteins interact with H3/H4 acetylated tails through bromodomains and may show specificity toward or against H3.3-containing nucleosomes (Adhikary et al., 2016; Guo et al., 2014; N. Li et al., 2016; Wen et al., 2014). Numerous studies have shown ZMYND8 interacts with and can recruit the SWI/SNF-like repressive NuRD (Mi2-β) chromatin remodeler complex (Adhikary et al., 2016; Gong et al., 2015; Savitsky et al., 2016; Spruijt et al., 2016), which contains HDACs and interacts with H3.3 (Kraushaar et al., 2018). Like ARID1A, ZMYND8 has also been shown to suppress super-enhancer hyperactivation (Shen et al., 2016), and, more recently, ZMYND8 and ARID1A were identified in the same screen as key chromatin regulators of EMT (Serresi et al., 2021). Therefore, we investigated the potential roles of ZMYND8 and possible co-factors as mediators of the observed ARID1A-H3.3 co-regulation (Figure 5B).

**Figure 5.**
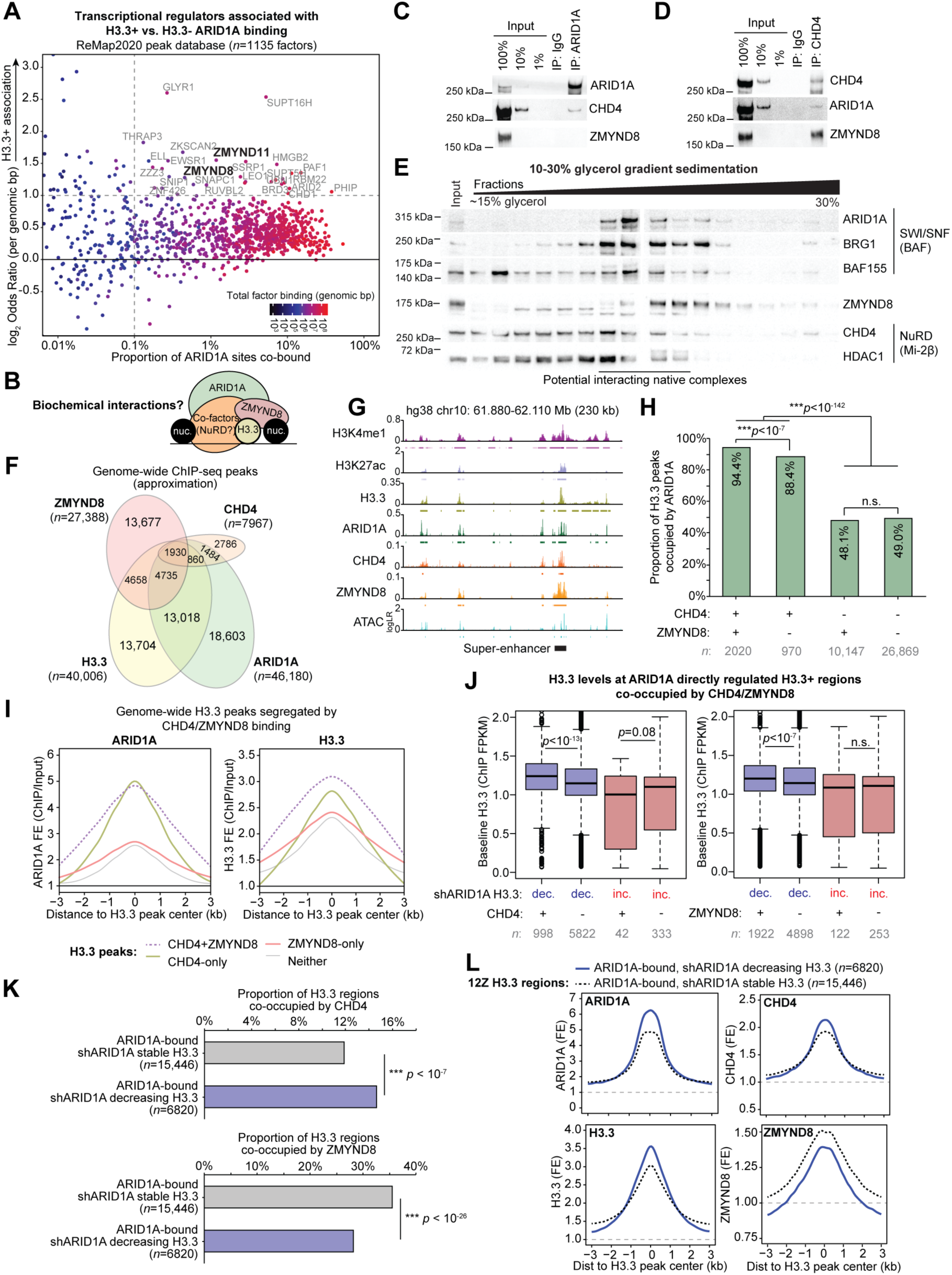
Biochemical and genomic characterization of ARID1A, CHD4, and ZMYND8 chromatin interactions co-regulating H3.3. **A**, Genome-wide associations between ARID1A binding at H3.3+ vs. H3.3-regions for all 1135 transcriptional regulator peak sets included in the ReMap2020 peak database. Labeled factors exhibit an H3.3+ ARID1A binding association with genomic odds ratio >2 and overlap with >0.1% of ARID1A binding sites. ZMYND11 and ZMYND8 (bolded) are two of the top factors most associated with H3.3+ ARID1A binding. **B**, Chromatin model schematic depicting hypothesized relationship between ARID1A-SWI/SNF and ZMYND8 co-regulation of H3.3, possibly mediated by co-factors. **C**, ARID1A co-immunoprecipitation detecting physical interaction with NuRD catalytic subunit CHD4, but not ZMYND8. **D**, CHD4 co-immunoprecipitation detecting physical interactions with both ARID1A and ZMYND8. **E**, 10-30% glycerol gradient sedimentation and immunoblotting for SWI/SNF, NuRD, and ZMYND8. Relative fractions display native protein complexes transitioning from low molecular weight (left) to high molecular weight (right). Underlined fractions highlight potential interacting native complexes containing ZMYND8 and members of SWI/SNF (BAF) and NuRD (Mi-2β). **F**, Genome-wide ChIP-seq peak overlaps between ARID1A, CHD4, ZMYND8, and H3.3. Peak numbers within the Euler diagram are approximations and not mutually exclusive due to varying peak sizes. **G**, Example locus on chromosome 10 displaying ARID1A, CHD4, ZMYND8, and H3.3 co-regulation. **H**, Enrichment for ARID1A co-regulation of H3.3 peaks bound by CHD4 and/or ZMYND8. Statistic is two-tailed Fisher’s exact test. **I**, Average ChIP-seq signal density histograms for ARID1A (left) and H3.3 (right) at H3.3 peaks bound by CHD4 and/or ZMYND8. **J**, H3.3 abundance (ChIP FPKM) at ARID1A-bound shARID1A differential H3.3 regions co-bound by CHD4 or ZMYND8. Statistic is two-tailed, unpaired Wilcoxon’s test. **K**, Positive association between CHD4 binding (top) and negative association between ZMYND8 binding (bottom) and ARID1A direct maintenance of H3.3 chromatin, genome-wide. Statistic is two-tailed Fisher’s exact test. **L**, Average ChIP-seq signal density histograms for ARID1A, H3.3, CHD4, and ZMYND8 across ARID1A-bound H3.3 regions that decreased or were stable with shARID1A.

ARID1A co-immunoprecipitation (co-IP) using an anti-ARID1A antibody was first used to detect physical nuclear interactions with ZMYND8. We previously confirmed the specificity of the anti-ARID1A antibody by co-immunoprecipitation followed by mass spectrometry (Wilson et al., 2019). While ZMYND8 was not detected in the ARID1A pulldown following high salt washes (300 mM KCl), the NuRD catalytic subunit CHD4 was evident (Figure 5C). We then hypothesized that CHD4-NuRD may serve as an interface between ARID1A and ZMYND8. A reciprocal CHD4 co-IP confirmed nuclear interactions with both ARID1A and ZMYND8 (Figure 5D). To further support that ARID1A, ZMYND8, and CHD4 are found in high molecular weight nuclear complexes of similar size, glycerol gradient sedimentation was performed. Native fractions were observed that included ZMYND8 and members of both SWI/SNF and NuRD (Figure 5E). These data suggest that physical interactions between ARID1A and CHD4 may regulate H3.3 chromatin with support from ZMYND8.

We then examined genome-wide chromatin regulation by CHD4 and ZMYND8 in relation to ARID1A and H3.3. Genome-wide binding profiles of CHD4 and ZMYND8 were measured by ChIP-seq. Roughly 2000 genomic regions were identified with H3.3 and all three chromatin regulators co-localized (Figure 5F-G). Across all H3.3 peaks genome-wide, ARID1A binding was most strongly enriched at ZMYND8-CHD4 co-bound sites compared to sites occupied by either CHD4 or ZMYND8 alone (Figure 5H). Notably, when CHD4 was absent, ARID1A binding at H3.3 sites did not correlate with the presence of ZMYND8 (Figure 5H). These data suggest that CHD4 may be primarily responsible for ARID1A recruitment to H3.3 chromatin. We further investigated ARID1A binding and H3.3 abundance across H3.3 peaks segregated by the presence of CHD4/ZMYND8. ARID1A binding was again strongest at H3.3 peaks co-bound by CHD4 as opposed to those without CHD4 (Figure 5I). H3.3 abundance was similarly highest at CHD4-bound peaks, although CHD4+ZMYND8 peaks showed the overall highest H3.3 levels (Figure 5I). With respect to H3.3 regions dependent on ARID1A chromatin interactions, we observed that baseline H3.3 levels were significantly higher at regions that decreased in H3.3 following ARID1A knockdown if they were co-occupied by CHD4 or ZMYND8, but this was not observed at regions that gained H3.3 following ARID1A loss (Figure 5J). Intriguingly, we observed that genome-wide H3.3 regions directly maintained by ARID1A chromatin interactions are associated with CHD4 but not ZMYND8 (Figure 5K). Moreover, genome-wide regions that lose H3.3 following ARID1A loss due to disrupted ARID1A chromatin interactions tend to have higher baseline levels of ARID1A, CHD4, and H3.3, but lower levels of ZMYND8 in comparison to stable H3.3 regions (Figure 5L). Altogether, these results suggest CHD4 and ZMYND8 are associated with ARID1A-H3.3 co-regulation.

### ARID1A-CHD4-ZMYND8 suppress hyperactivation of H3.3-marked super-enhancers

Genome-wide analyses indicate that CHD4 and ZMYND8 co-regulation of ARID1A and H3.3 may be chromatin context-specific and occur at subsets of regulatory regions bound by ARID1A-CHD4-ZMYND8. While ARID1A loss causes widespread H3.3 reduction in chromatin, we observed that ARID1A-H3.3 co-regulation is most frequent at active enhancer and super-enhancer chromatin states. As such, we next examined ARID1A-CHD4-ZMYND8 co-regulation of H3.3 at enhancers. At ARID1A-bound active enhancers, defined as accessible (ATAC+) H3K27ac peaks located >3 kb from an annotated TSS (Figure 6A), ZMYND8 binding is associated with presence of CHD4, as expected (Figure 6B). Moreover, ZMYND8 binding was detected at 84.6% of ARID1A+CHD4 super-enhancers (*n* = 507) as opposed to 63.2% of ARID1A+CHD4 typical enhancers (*n* = 2282) (Figure 6B). Importantly, ARID1A, CHD4, and ZMYND8 co-binding at active enhancers is associated with presence of H3.3, and this association is greater at super-enhancers (Figure 6C). We previously observed that ARID1A suppresses H3K27-hyperacetylation at a subset of active super-enhancers (Wilson et al., 2020). ARID1A and CHD4 binding levels are not substantially different at suppressed super-enhancers that become hyperacetylated following ARID1A loss vs. those with stable acetylation (Figure 6D). Strikingly, ZMYND8 binding and H3.3 abundance are significantly higher at suppressed super-enhancers that become hyperacetylated (Figure 6D). These data indicate that ZMYND8 is associated with CHD4 and ARID1A most frequently at active H3.3-marked super-enhancers that are suppressed by ARID1A.

**Figure 6.**
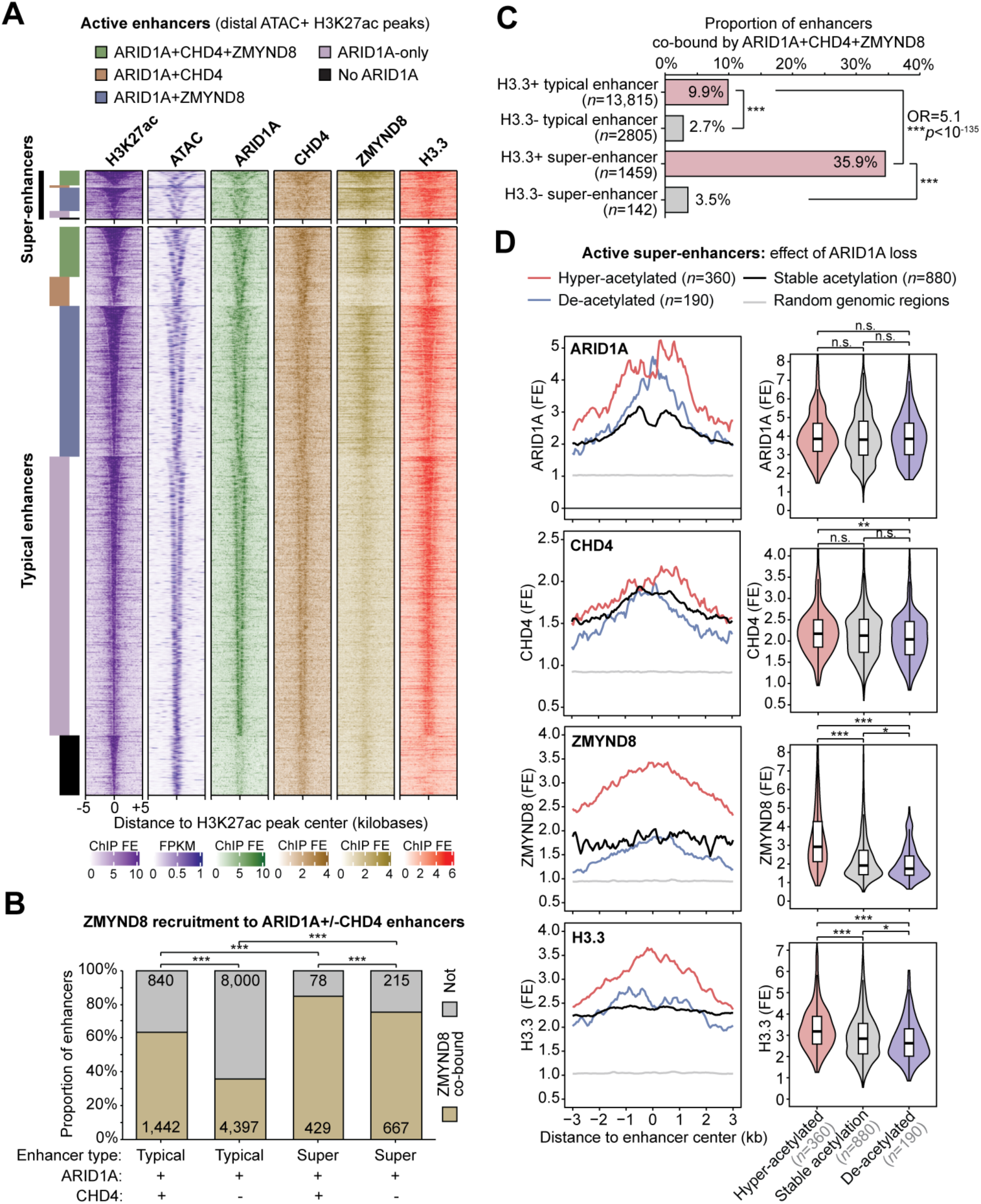
H3.3 enhancer regulation by ARID1A, CHD4, and ZMYND8. **A**, Heatmap of chromatin features at 15,925 active typical enhancers and 1374 distal super-enhancer constituents (H3K27ac peaks co-marked by ATAC) segregated by ARID1A ± CHD4 ± ZMYND8 binding. Enhancers are centered on the H3K27ac peak, and signal is displayed as indicated for the flanking 5 kb in either direction. **B**, ZMYND8 binding detection at ARID1A-bound typical and super-enhancers with or without CHD4 co-binding. Statistic is two-tailed Fisher’s exact test. **C**, Association between ARID1A+CHD4+ZMYND8 co-binding at enhancers and presence of H3.3. H3.3+ super-enhancers show the most frequent co-binding. Statistic is two-tailed Fisher’s exact test. **D**, Chromatin features at active super-enhancer constituents segregated by ARID1A loss-driven H3K27-acetylation dynamics: hyper-acetylated, de-acetylated, or stably acetylated. Left, average ChIP-seq signal density histograms across enhancer classes. Right, violin plots quantifying signal (ChIP/input fold-enrichment) across enhancer classes. Statistic is two-tailed, unpaired Wilcoxon’s test.

### ZMYND8 specifies ARID1A-CHD4 chromatin repression toward H4K16ac

Our data indicate the ZMYND8 module appears to be associated with repressive chromatin targeting by ARID1A at H3.3-marked super-enhancers. Histone tail reader functions of ZMYND8 are a plausible mechanism through its BRD, PWWP, and PHD domains, which interact with acetylated H3/H4 residues and methylated H3 residues (Savitsky et al., 2016). Particularly, the ZMYND8 bromodomain was recently described to interact with acetylated H4 tails and recruit CHD4 to repress transcription following DNA damage (Gong et al., 2015). However, NuRD also interacts with other histone substrates such as H2A.Z, which has been linked to H3.3-mediated transcriptional poising (Chen et al., 2013). Of further relevance, P300 was recently shown to acetylate H2A.Z when stimulated by recognition of H4 acetylated residues through its bromodomain (Colino-Sanguino et al., 2019). To better resolve our model of ARID1A-CHD4-ZMYND8-mediated H3.3 chromatin repression, we generated a more comprehensive genome-wide chromatin state model that contains 5 additional features related to reported ZMYND8 and NuRD activity that were profiled both before and after ARID1A knockdown: H3.3, pan-acetyl-H4 (K5/K8/K12/K16, pan-H4ac), H4K16ac, H2A.Z, and acetyl-H2A.Z (K4/K7; H2A.Zac) (Figure 7A, Figure 7—figure supplement 1). Since the anti-H2A.Zac antibody used in our assays has not been characterized for specificity towards acetylated H2A.Z containing peptides, we performed a histone peptide array (Rothbart, Krajewski, Strahl, & Fuchs, 2012) and found that it specifically recognizes H2A.Zac (Figure 7—figure supplement 2). The new 12-feature chromatin state model was quantitatively optimized at 25 chromatin states (see Materials and Methods for details) (Figure 7—figure supplement 3). The enhanced resolution of our model revealed new state identities such as H3.3+ poised bivalent promoters (state 8), H2A.Zac+ poised enhancers (state 14), and notably the segregation of upstream active promoter super-enhancers into H4(K16)ac+ (state 2) and negative (state 5) classes (Figure 7A). We next determined states enriched for co-binding of ARID1A-CHD4-ZMYND8 and shARID1A direct decreasing H3.3 (Figure 7B). Both upstream active promoter super-enhancer states (states 2 and 5) showed the strongest enrichment for shARID1A direct decreasing H3.3, while ZMYND8 binding and ARID1A-CHD4-ZMYND8 co-binding was most enriched at the H4(K16)ac+ upstream active promoter super-enhancer class (state 2) (Figure 7B).

**Figure 7.**
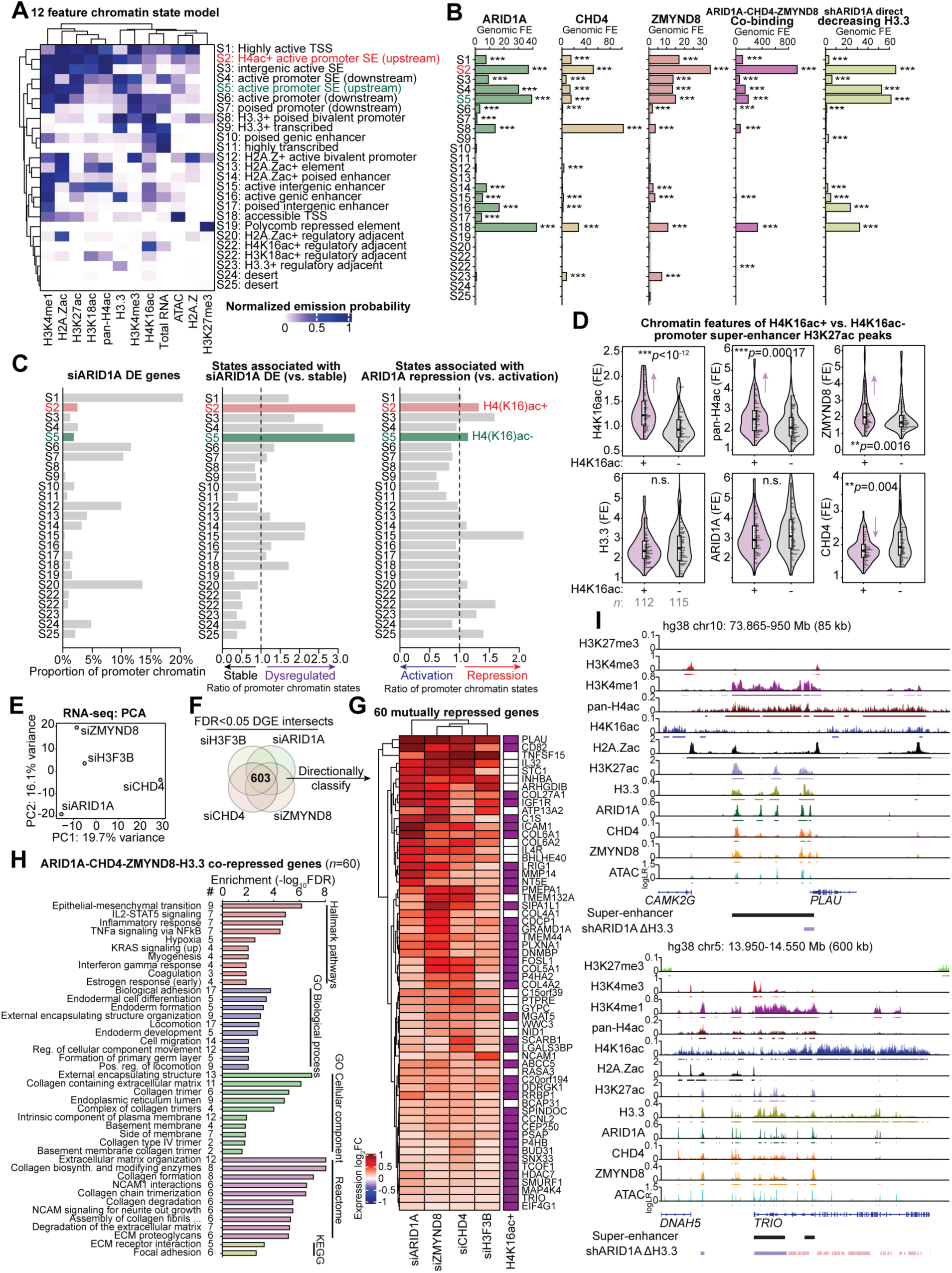
ZMYND8-mediated chromatin repression is specified by H4(K16)ac. **A**, Heatmap of clustered, normalized feature emission probabilities and associated functional annotation of the new 12 feature, genome-wide chromatin 25-state model. States (S_) are labeled based on order of normalized emission probability clustering. See methods for details on optimal model selection. **B**, Genomic fold-enrichment (FE) for ARID1A, CHD4, ZMYND8, co-binding, and shARID1A direct decreasing H3.3 among the 25 chromatin states. Statistic is hypergeometric enrichment test. **C**, Modeled chromatin states among reference gene promoter regions (±3 kb around annotated TSS). Left, proportion of promoter chromatin for siARID1A DE genes (*DESeq2*, FDR < 0.0001) belonging to each of the 25 states. Center, ratio of promoter chromatin states associated with siARID1A DE genes (FDR < 0.0001) compared to stable genes (FDR > 0.05). Right, ratio of promoter chromatin states associated with ARID1A transcriptional repression (i.e. siARID1A upregulation) compared to activation (i.e. siARID1A downregulation). **D**, Violin plots quantifying chromatin feature signal at H4K16ac+ (purple) vs. H4K16ac- (gray) promoter super-enhancer constituent H3K27ac peaks. Statistic is two-tailed, unpaired Wilcoxon’s test. **E**, Principal component analysis (PCA) of RNA-seq expression log_2_FC (shrinkage-corrected) values for siCHD4, siZMYND8, siH3F3B, and siARID1A treatment conditions vs. controls. 1974 genes with _s_log_2_FC variance >0.1 were used for PCA. **F**, Schematic of identifying mechanistic genes co-repressed by ARID1A, H3.3, CHD4, and ZMYND8, i.e., upregulated (*DESeq2*, FDR < 0.05) with siARID1A, siH3F3B, siCHD4, and siZMYND8 treatments. **G**, Clustered heatmap of expression log_2_FC values for 60 co-repressed genes upregulated in all 4 knockdown conditions. Rightmost column demarcates presence of H4K16ac peaks over promoter or gene body. **H**, Top gene sets enriched (hypergeometric enrichment test, FDR < 0.05) among the 60 ARID1A-CHD4-ZMYND8-H3.3 co-repressed genes from various gene set databases. **I**, Example target gene loci, *PLAU* and *TRIO*, marked by nearby H3.3+ super-enhancers within H4(K16)ac+ domains that are co-bound by ARID1A, CHD4, ZMYND8, where ARID1A loss leads to significant depletion of H3.3 (ChIP-seq FDR < 0.05), and knockdown of ARID1A, H3.3, CHD4, or ZMYND8 leads to significant expression upregulation (RNA-seq FDR < 0.05).

We further investigated chromatin state identities at reference annotated gene promoter regions (±3 kilobases flanking TSS). As expected, the highly active TSS state (state S1) is the most prevalent promoter chromatin state identity at genes transcriptionally regulated by ARID1A (siARID1A DE genes), while upstream active promoter super-enhancer states (states 2 and 5) are relatively rare (Figure 7C, left). However, comparing the promoter chromatin state identities across siARID1A DE vs. stable genes revealed that upstream active promoter super-enhancer states are associated with ARID1A transcriptional regulation (Figure 7C, center). We further segregated promoter chromatin based on genes that are upregulated with siARID1A (i.e. repressed by ARID1A) vs. downregulated with siARID1A (i.e. activated by ARID1A). The H4(K16)ac+ active promoter super-enhancer state marked by high ARID1A-CHD4-ZMYND8 co-binding (state 2) showed stronger enrichment for ARID1A transcriptional repression than those without H4(K16)ac (state 5) (Figure 7C, right). In agreement, we also observed that chromatin accessibility directly repressed by ARID1A is associated with presence of H4 acetylation (Figure 7—figure supplement 4). Promoter super-enhancer H3K27ac peaks were next directly segregated by detection of H4K16ac (H4K16ac+, *n* = 112; H4K16ac-, *n* = 115). ARID1A binding and H3.3 abundance were not significantly different between H4K16ac stratified super-enhancers, but ZMYND8 binding was stronger at the H4K16ac+ regions, while CHD4 binding was lower at H4K16ac+ regions (Figure 7D). Further, a correlation of measured chromatin features across all 227 promoter-proximal super-enhancer constituent enhancers supported that ZMYND8 binding is associated with acetylated H4 marks (Figure 7—figure supplement 5). These analyses collectively suggest that H4(K16)ac marked promoter super-enhancers may recruit repressive ARID1A-CHD4-ZMYND8 complexes to regulate H3.3.

To identify genes targeted by the ARID1A-CHD4-ZMYND8 regulation of repressive H3.3, we also used siRNA to deplete CHD4 (siCHD4) and ZMYND8 (siZMYND8) followed by RNA-seq (Figure 7E, Figure 7—figure supplement 6A-F). As expected, we observed enrichment of expression alterations following loss of ZMYND8 and CHD4 among genes with detected promoter binding by each factor (Figure 7—figure supplement 6G-H). To further support whether ZMYND8 is associated with transcriptional repression by ARID1A and H3.3, we investigated ZMYND8-mediated gene expression at siARID1A/siH3F3B DGE directional classes (refer to Figure 4G-H). Indeed, siZMYND8 gene expression alterations were strongly enriched at genes mutually repressed by ARID1A and H3.3 compared to other gene classes, where ZMYND8 also functioned mostly as a repressor (Figure 7—figure supplement 6I).

Differential gene expression again revealed highly overlapping genes transcriptionally regulated by ARID1A, CHD4, ZMYND8, and H3.3 (Figure 7—figure supplement 6J-L), including 603 genes affected by each of the four knockdowns (FDR < 0.05) (Figure 7F, Figure 7—figure supplement 6L). These included 60 genes mutually repressed by ARID1A, CHD4, ZMYND8, and H3.3 (Figure 7G). These mechanistic co-repressed genes were enriched for EMT, adhesion, development, locomotion, collagens, and extracellular matrix gene sets (Figure 7H). Further, 68% of these genes were marked by gene body H4K16ac, an enrichment compared to less than half of all expressed genes (Figure 7—figure supplement 7). Two physiologically relevant target genes revealed through integrative epigenomic analysis are *PLAU* and *TRIO*, both of which are located within broad H4K16ac+ domains and near active H3.3+ super-enhancers co-bound by ARID1A, CHD4, and ZMYND8 (Figure 7I). ARID1A loss leads to decreased promoter H3.3 abundance and transcriptional hyperactivation of *PLAU* and *TRIO* (Figure 7I). We also observed that co-knockdown of ARID1A and CHD4 led to increased induction of *PLAU* compared to either knockdown separately (Figure 7—figure supplement 8).

### ARID1A-H3.3 repressed chromatin targets are aberrantly activated in human endometriomas

Our studies in the 12Z human endometrial epithelial cell line have revealed a mechanism of cooperative chromatin regulation by ARID1A-CHD4-ZMYND8 maintenance of repressive H3.3. To support the relevance of these chromatin regulatory networks on pathologically related gene expression, we utilized a transcriptome expression data set comparing human endometriomas to control endometrial tissue samples (Hawkins et al., 2011). Endometriomas are a result of ectopic spread of endometrial tissue onto the ovary, forming cysts associated with ovarian cancer development (Grandi et al., 2015; Zondervan et al., 2020). Three ARID1A-H3.3 related gene sets were investigated for relevance in human endometrioma gene expression alterations: 1) shARID1A direct decreasing promoter H3.3 genes (*n* = 412), 2) ARID1A-H3.3 co-repressed genes (i.e. siARID1A/siH3F3B upregulated, FDR < 0.001, *n* = 196), and 3) ARID1A-H3.3-CHD4-ZMYND8 co-repressed genes (i.e. upregulated with any knockdown, FDR < 0.05, *n* = 60). We observed significant enrichment for all three of these gene sets among human endometrioma DGE (Figure 8A, left). Moreover, the overlapping DE genes were more likely to be upregulated in endometriomas than expected by chance, indicating relief of repression is also observed in pathology (Figure 8A, right). Similarly, examining the endometrioma vs. control endometrium expression log_2_FC values indicated that each gene set tended to be overall upregulated in the pathological, pre-cancerous state (Figure 8B). Mechanistic genes aberrantly activated in endometriomas that could be attributed to disruption of ARID1A-H3.3 chromatin repression mechanisms include *C1S*, *SCARB1*, *GYPC*, *WWC3*, *COL6A2*, and *MAP4K4* (Figure 8C). Our data indicate that ARID1A cooperatively maintains transcriptionally repressive H3.3 chromatin through protein-protein interactions with CHD4 and ZMYND8, likely recruited to H4(K16)ac+ chromatin, such that loss of any of these factors leads to alleviation of H3.3-mediated repression and consequential aberrant gene activation in various endometrial disease contexts where ARID1A mutations are thought to drive pathogenesis.

**Figure 8.**
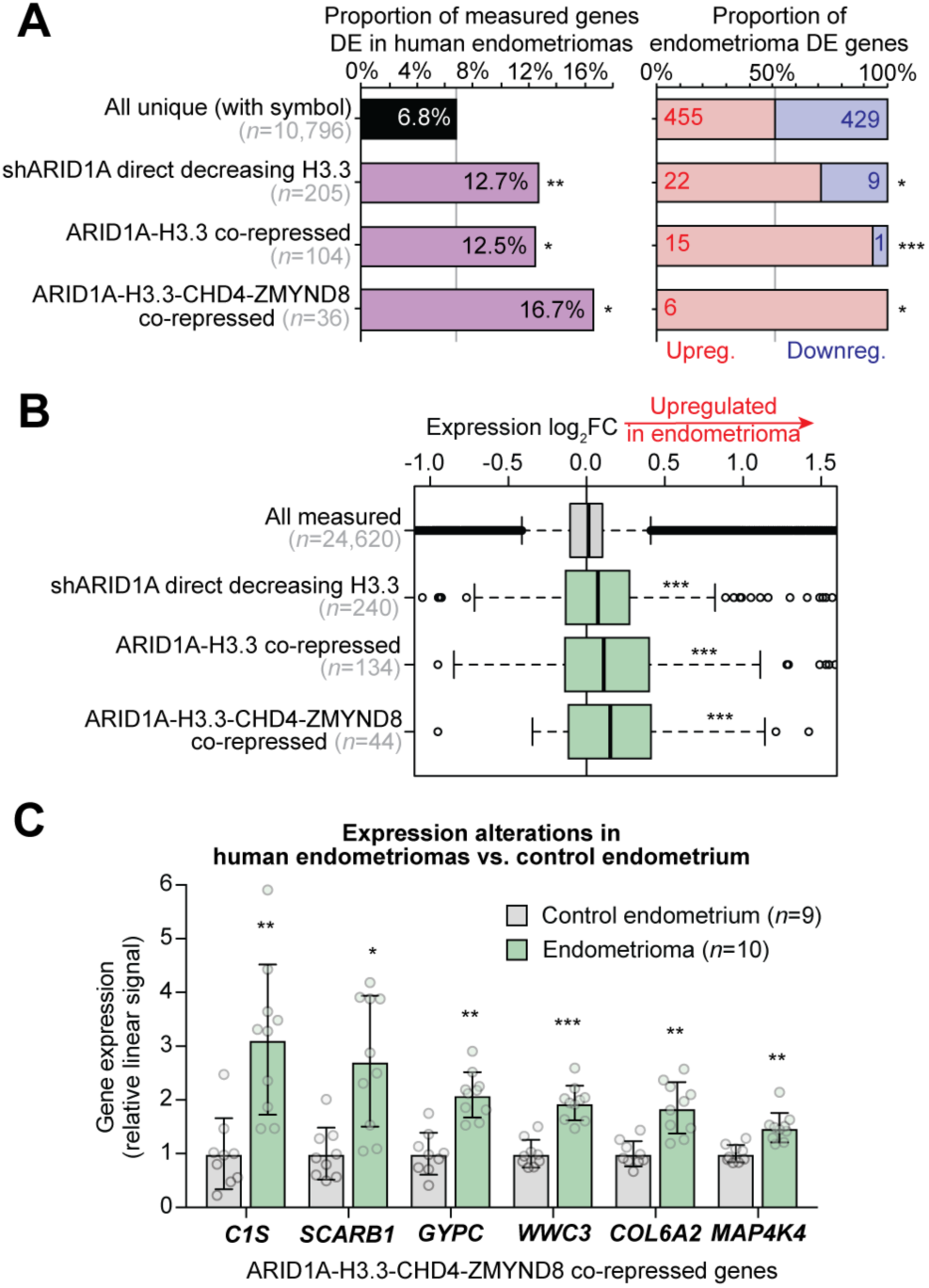
Mechanistic gene expression alterations in human endometriomas. **A**, Left, enrichment for ARID1A-H3.3 co-repressive chromatin mechanistic gene sets among human endometrioma (ovarian endometriosis) vs. control endometrium DE genes reported by Hawkins et al. (Hawkins et al., 2011), compared to all unique measured genes. Right, proportion of overlapping DE genes that are upregulated vs. downregulated in endometriomas, compared to all unique measured genes. Statistic is hypergeometric enrichment. **B**, Box plots displaying endometrioma expression log_2_FC values for probes annotated to genes within mechanistic gene sets, compared to all measured probes. Statistic is two-tailed, unpaired Wilcoxon’s test. **C**, Relative expression box-dot plots of 6 genes upregulated in endometriomas vs. control endometrium that are co-repressed by ARID1A, H3.3, CHD4, and ZMYND8. Statistic is *limma* FDR-adjusted *p*. * *p* < 0.05, ** *p* < 0.01, *** *p* < 0.001.

## Discussion

We have provided evidence that ARID1A functions to maintain the variant histone H3.3 in active chromatin. ARID1A loss leads to H3.3 depletion at active enhancers and super-enhancers, due to disrupted ARID1A chromatin interactions, leading to H3.3 redistribution toward active genic and transcribed elements. SWI/SNF functions to eject nucleosomes and open chromatin, rather than assemble nucleosomes on chromatin. SWI/SNF disruption of nucleosomes may be required for H3.3 incorporation and thus be coupled to nucleosome assembly. Therefore, we hypothesize that H3.3 regulation by ARID1A-SWI/SNF occurs by ejecting canonical H3.1 containing nucleosomes in favor of H3.3 incorporation by other assembly or chaperone factors. In the absence of ARID1A-SWI/SNF, H3.3 nucleosome assembly may be impeded. Given previous associations between H3.3 epigenetic memory and cell fate plasticity, it is intriguing to consider a role for ARID1A-SWI/SNF regulation of H3.3 as being a critical determinant of endometrial epithelial cell identity and homeostasis across the menstrual cycle when proliferation and differentiation occur (Szenker et al., 2011).

ARID1A maintenance of H3.3 is associated genomic interactions with CHD4, a catalytic subunit in the SWI/SNF-like NuRD remodeler complex. Target regulatory activity is further directed by CHD4 interactions with histone reader ZMYND8, where ZMYND8 is recruited to H4(K16)ac-marked chromatin, possibly through its bromodomain. This ZMYND8-mediated targeting of ARID1A-CHD4 to H3.3 is associated with repressive chromatin regulation, notably at super-enhancers located upstream of genes, such that disruption of this chromatin mechanism causes relief of repression and subsequent transcriptional hyperactivation. Plasminogen activator urokinase (*PLAU*) was identified as a key target gene repressed by this mechanism in our 12Z endometriotic epithelial cell model. *PLAU* was also recently observed as transcriptionally activated during human menstruation (Wang et al., 2020), suggesting similar repressive chromatin mechanisms may govern *PLAU* regulation in the healthy endometrium. *PLAU* is upregulated in ovarian endometrioid carcinomas from women with concurrent endometriosis (C. Zhang et al., 2018), suggesting *PLAU* upregulation may promote malignant transformation in endometriosis. ARID1A mutations are frequently observed in endometriosis-associated ovarian cancers (Jones et al., 2010; Wiegand et al., 2010). *C1S*, a component of the complement C1 complex, is another gene that is transcriptionally repressed by ARID1A, CHD4, ZMYND8, and H3.3 that is aberrantly upregulated in human endometriomas. It has been reported that the complement system is activated in women with endometriosis (Sikora et al., 2018), suggesting that ARID1A mutation and associated disruption of chromatin repression may be a possible disease mechanism.

H3.3 is considered an active chromatin mark associated with transcriptional activation. However, our data and others have demonstrated that H3.3 can play roles in transcriptional repression, as well as transcriptional poising and higher-order chromatin regulation, although the mechanisms governing these functional specificities remain unclear (Shi et al., 2017). A simple hypothesized mechanism explaining how H3.3 can function repressively is through associations with CHD4 and the NuRD complex, as we have studied here. Historically, NuRD has been studied as a repressor due to its subunit composition that includes the histone deacetylases HDAC1/2, although activating roles of NuRD are also known (Basta & Rauchman, 2015; Tong, Hassig, Schnitzler, Kingston, & Schreiber, 1998; Xue et al., 1998). An early study of H3.3 chromatin dynamics indicated that NuRD components were associated with active regions marked by high H3.3 turnover (Ha, Kraushaar, & Zhao, 2014). More recently, NuRD has been shown to directly interact with H3.3 nucleosomes (Kraushaar et al., 2018). Further, we have shown that ARID1A physically interacts with CHD4 in endometrial epithelial cells, and ARID1A and CHD4 co-binding at unique chromatin sites is associated with H3.3 maintenance and transcriptional repression.

Despite these observations, it is not clear how ARID1A cooperates with CHD4 through protein-protein interactions to regulate H3.3 nucleosomes. Intriguingly, CHD4/NuRD was recently shown to control super-enhancer accessibility and maintain lower acetylation levels through its HDAC activity (Marques et al., 2020), similar to our findings with ARID1A by antagonizing P300 (Wilson et al., 2020). In support of our data, the authors observed CHD4 interactions with SWI/SNF. Recently, NuRD and SWI/SNF recruitment to active TSS and enhancers was impaired in H3.3K4A mutant mouse ESCs (Gehre et al., 2020), suggesting that NuRD and SWI/SNF recruitment is dependent on the K4 residues on H3.3. The same study also reported the K4A mutation destabilized H3.3 and that both CHD4 and SMARCA4 (SWI/SNF) are required for H3.3 stability, further supporting a role for NuRD and SWI/SNF in H3.3 maintenance (Gehre et al., 2020). Loss of H3.3 following disruption of ARID1A chromatin interactions could lead to destabilization of repressive NuRD. One hypothesized cooperative mechanism that could explain how ARID1A and CHD4 maintain H3.3 in active chromatin is through different aspects of remodeler activity conferred by each complex. ARID1A could remodel or remove canonical H3 nucleosomes—a typical role of SWI/SNF—while CHD4 could function to assemble or stabilize H3.3. In this scenario, failure to remove canonical H3 following ARID1A loss may impair the ability of CHD4 to assemble or stabilize H3.3, and thus loss of either ARID1A or CHD4 may break this cycle. The related CHD1 remodeler is known to be required for H3.3 deposition into chromatin *in vivo* (Konev et al., 2007), further suggesting a necessary role for remodeler activity in H3.3 nucleosome assembly. It is important to note that H3.3 has a defined chaperone associated with transcriptional activity, HIRA (Shi et al., 2017). P400 is another SWI/SNF-like remodeler recently shown to exchange H3.3 nucleosomes that could also possibly collaborate with SWI/SNF (Pradhan et al., 2016).

*In silico* analyses from the ReMap 2020 transcriptional regulator peak database (Cheneby et al., 2020) predicted that ZMYND8 is highly associated with H3.3 chromatin regulation by ARID1A. While we did not detect physical interactions directly between ARID1A and ZMYND8 under high salt conditions, we did observe ARID1A-CHD4 and CHD4-ZMYND8 interactions. ZMYND8 has been shown to interact with NuRD in numerous contexts (Adhikary et al., 2016; Gong et al., 2015; Savitsky et al., 2016; Spruijt et al., 2016). Intriguingly, one recent study reported that ZMYND8 directly recognizes mutant H3.3G34R (Jiao et al., 2020). Our data indicate that ZMYND8 links repressive H3.3 to H4 acetylation. In support, the ZMYND8 bromodomain directly interacts with acetylated H4 tails (Adhikary et al., 2016), and TIP60-mediated H4 acetylation can functionally recruit ZMYND8 through this mechanism to repress transcription with CHD4 in response to DNA damage (Gong et al., 2015). H4K16ac was also previously reported to be required for the NoRC repressor complex to bind and silence chromatin (Zhou & Grummt, 2005). Our data also indicate that ARID1A directly suppresses chromatin accessibility at sites marked by H4 acetylation, suggesting that SWI/SNF chromatin remodeler activity may be involved in ZMYND8-NuRD-mediated chromatin repression. Interestingly, in addition to NuRD-dependent regulation, ZMYND8 regulates transcription independently of NuRD (Adhikary et al., 2016). Positive transcriptional regulation by ZMYND8 has been shown to occur through its association with the P-TEFb elongation complex (Ghosh et al., 2018). ZMYND8-NuRD repression in response to DNA damage was previously shown to rely on KDM5A demethylase activity (Gong, Clouaire, Aguirrebengoa, Legube, & Miller, 2017), further suggesting other factors may orchestrate this regulatory activity. ZMYND8 has been reported to be a super-enhancer factor that suppresses hyperactivation (Shen et al., 2016). Corroborating our results, ZMYND8 was previously shown to associate with NuRD at super-enhancers (Spruijt et al., 2016). We found that super-enhancers that become hyperacetylated following ARID1A loss are normally associated with the highest levels of H3.3 and ZMYND8 binding. In our proposed model, ZMYND8 bromodomain interactions with H4 acetylated tails recruit CHD4 and ARID1A for transcriptional repression at active chromatin. This most notably occurs at H3.3+ super-enhancers, where all three factors co-localize most frequently. Further work will seek to explain how ZMYND8 specifies this repressive activity, as CHD4 and ARID1A can also function toward transcriptional activation.

It is worth considering alternative hypotheses of how transcriptional homeostasis could be altered following disruption of these chromatin regulators and epigenomic marks. In addition to promoter and enhancer chromatin regulation, SWI/SNF, NuRD, and ZMYND8 have been shown to mediate transcriptional pausing and elongation by Pol II and associated machinery (Bottardi et al., 2014; Ghosh et al., 2018; Schwabish & Struhl, 2007; Trizzino et al., 2018), as well as DNA repair (Gong et al., 2015; Park et al., 2006; Smeenk et al., 2010). Super-enhancers mark critical cell identity genes (Whyte et al., 2013), and recent evidence suggests chromatin mechanisms coupling transcription and DNA repair occur at super-enhancers to control transcriptional hyperactivation (Hazan, Monin, Bouwman, Crosetto, & Aqeilan, 2019). Super-enhancer chromatin co-regulation by ARID1A, CHD4, and ZMYND8 may fine tune transcriptional activation states and thus reflect a mechanism at the intersection of transcriptional regulation and DNA repair.

## Materials and Methods

### Data availability

All new data generated in this study have been deposited to the Gene Expression Omnibus (GEO) at SuperSeries accession GSE190557, and access is available with reviewer token: ijmtkuuivtwzbwz. Previously generated and re-analyzed data were retrieved from GEO at accessions GSE121198 and GSE148474 and analyzed as previously described (Wilson et al., 2020; Wilson et al., 2019).

### Cell culture, siRNA transfections, and lentiviral shRNA particle usage

Adherent, human 12Z endometriotic epithelial cells were cultured in DMEM/F12 media in the presence of 10% serum (FBS), 1% L-glutamine, and 1% penicillin/streptomycin. Cells were seeded in antibiotic-free media the day before siRNA transfection. 50 nM siRNA (Dharmacon, ON-TARGETplus) were transfected into cells using the Lipofectamine RNAiMAX (ThermoFisher Scientific) reagent, according to the manufacturer protocol, in OptiMEM (Gibco). Growth media was replaced 24 hours following transfection, without antibiotics. 48 hours after transfection, low serum (0.5% FBS) growth media was added with antibiotics. Cells were harvested 72 hours following siRNA transfection. Lentiviral shRNA particles were prepared with Lenti-X 293T cells (Takara) and MISSION pLKO.1 plasmids (Sigma-Aldrich) as previously described (Wilson et al., 2020). Lentiviral shRNA particles were titered using the qPCR Lentiviral Titration Kit (ABM). shRNA particles were transduced into 12Z cells at a 100-fold multiplicity of infection, and media was replaced 24 hours later. Cells were harvested 72 hours following shRNA transduction.

### Cell cycle analysis

The Click-iT Plus EdU Flow cytometry Assay Kit (Invitrogen) was used for cell cycle assays. 12Z cells were treated with 10 mM of EdU for 2 hours in culture media. Cells were harvested by trypsinization and washed in 1% BSA in PBS. Cells were resuspended in 100 µL of ice-cold PBS, and 900 µL of ice cold 70% ethanol was added dropwise while vortexing. Cells were incubated on ice for two hours. Cells were washed with 1% BSA in PBS and then treated with the Click-iT Plus reaction cocktail including Alexa Fluor 488 picolyl azide according to the manufacturer’s instructions for 30 minutes. Cells were washed with 1X Click-iT permeabilization buffer and wash reagent, and then treated with 5 mM of Vybrant DyeCycle Ruby Stain (ThermoFisher) diluted in 1% BSA in PBS for 30 minutes at 37 °C. Flow cytometry was performed using a BD Accuri C6 flow cytometer (BD Biosciences) and analyzed using FlowJo v10 software (BD Biosciences).

### Histone extraction

12Z cells were washed with PBS and scraped in PBS containing 5 mM sodium butyrate. Cells were centrifuged and resuspended in TEB buffer (PBS supplemented with 0.5% Triton X-100, 5 mM sodium butyrate, 2 mM phenylmethylsulfonyl fluoride, 1X protease inhibitor cocktail) and incubated on a 3D spindle nutator at 4 °C for 10 minutes. Cells were centrifuged at 3000 RPM for 10 minutes at 4 °C. TEB wash step was repeated once. Following second wash, pellet was resuspended in 0.2 N HCl, and incubated on 3D spindle nutator at 4 °C overnight. The following day, samples were neutralized with 1:10 volume 1M Tris-HCl pH 8.3. Sample was centrifuged at 3000 RPM for 10 minutes at 4 °C, and supernatant containing histone proteins was collected.

### Co-immunoprecipitation (co-IP)

Nuclear extracts were prepared as previously described (Chandler et al., 2013), dialyzed overnight into 0% glycerol (25 mM HEPES, 0.1 mM EDTA, 12.5 mM MgCl2, 100 mM KCl, 1 mM DTT) using a Slide-A-Lyzer G2 Dialysis Cassette (10 kDa cutoff, ThermoFisher Scientific), and quantified with the BCA Protein Assay Kit (Pierce, ThermoFisher Scientific). Primary antibodies (anti-ARID1A, D2A8U, Cell Signaling; anti-CHD4, D8B12, Cell Signaling;) were conjugated to Protein A Dynabeads (Invitrogen) overnight at 4 °C in 1X PBS + 0.5% BSA. Normal rabbit IgG (Cell Signaling) IPs were performed in parallel at equivalent masses, as negative controls. 500 µg nuclear lyase was diluted into IP buffer (20 mM HEPES, 150 mM KCl, 10% glycerol, 0.2 mM EDTA, 0.1% Tween-20, 0.5 mM DTT) to a final volume of 1 mL and clarified by centrifugation. After overnight IP at 4 °C, bead slurries were washed with a series of IP buffers at different KCl concentrations: 2X washes at 150 mM, 3X washes at 300 mM, 2X washes at 100 mM, 1X wash at 60 mM. Immunoprecipitants were eluted in 2X Laemmli buffer + 100 mM DTT at 70 °C for 10 minutes with agitation.

### Glycerol gradient sedimentation

Nuclear extracts were prepared, dialyzed, and quantified as described in the co-IP methods section. Density sedimentation by glycerol gradient was performed and probed similar to published reports (Mashtalir et al., 2018). Briefly, 4.5 mL 10-30% linear glycerol gradients were prepared using an ÄKTA start (Cytiva) from density sedimentation buffer (25 mM HEPES, 0.1 mM EDTA, 12.5 mM MgCl2, 100 mM KCl, 1 mM DTT) additionally containing 30% and 10% glycerol for initial and target concentrations, respectively. 200 µg nuclear lyase was overlaid on the glycerol gradient followed by ultracentrifugation at 40,000 rpm in an AH-650 swinging bucket rotor (ThermoFisher Scientific) for 16 hours at 4 °C. 225 µL gradient fractions were collected and concentrated using StrataClean resin (Agilent). Concentrated fractions were eluted in 1.5X Laemmli buffer + 37.5 mM DTT and run on SDS-PAGE for immunoblotting.

### Immunoblotting

Whole-cell protein lysates were prepared as previously described (Wilson et al., 2020). Proteins were quantified with the BCA Protein Assay Kit (Pierce, ThermoFisher Scientific). Protein samples in Laemmli buffer + DTT were denatured at 94 °C for 3 minutes prior to running on SDS-PAGE gels (6% gels for co-IP and glycerol gradients, 15% gels for histone extracts, and 4-15% gradient gels for whole-cell protein lysates). Gels containing histone extracts were wet transferred to nitrocellulose membranes at 4 °C for 3 hours at 400 mA current, then dried at room temperature followed by re-hydration in TBS + 0.1% Tween-20 (TBS-T) and blocking with Odyssey blocking buffer (LI-COR). All other gels were semi-dry transferred to PVDF using a Trans-Blot Turbo (Bio-Rad) according to the manufacturer’s protocol designed for high-molecular weight proteins, and blocked with either 5% BSA or 5% milk in TBS. The following primary antibodies were used: anti-ARID1A (D2A8U, Cell Signaling), anti-CHD4 (D4B7, Cell Signaling), anti-ZMYND8 (A302-089, Bethyl), anti-ZMYND8 (Atlas), anti-BRG1 (ab110641, abcam), anti-BAF155 (D7F8S, Cell Signaling), anti-HDAC1 (10E2, Cell Signaling), anti-histone H3.3 (ab176840, abcam), anti-histone H3.3 (2D7-H1, abnova), and anti-histone H3 (D1H2, Cell Signaling). IRDye fluorescent dye (LI-COR) secondary antibodies were used for LI-COR fluorescence-based protein visualization of histones. Horseradish peroxidase (HRP) conjugated secondary antibodies (Cell Signaling) were used for chemiluminescence-based protein visualization of all other targets. Clarity Western ECL substrate (Bio-Rad) was used to activate HRP for chemiluminescence, captured by ChemiDoc XRS+ imaging system (Bio-Rad).

### mRNA-seq and analysis

72 hours after initial siRNA transfection, and 24 hours after low-sera conditioning, 12Z cells were purified for RNA using the Quick-RNA Miniprep Kit (Zymo Research). Transcriptome libraries (n = 3 replicates) were prepared and sequenced by the Van Andel Genomics Core from 500 ng of total RNA using the KAPA mRNA HyperPrep kit (v4.17) (Kapa Biosystems). RNA was sheared to 300-400 bp. Prior to PCR amplification, cDNA fragments were ligated to IDT for Illumina unique dual adapters (IDT DNA Inc). Quality and quantity of the finished libraries were assessed using a combination of Agilent DNA High Sensitivity chip (Agilent Technologies), QuantiFluor dsDNA System (Promega), and Kapa Illumina Library Quantification qPCR assays (Kapa Biosystems). Individually indexed libraries were pooled and 50 bp, paired-end sequencing was performed on an Illumina NovaSeq 6000 sequencer using a 100-cycle sequencing kit (Illumina). Each library was sequenced to an average raw depth of 20-25 million reads. Base calling was done by Illumina RTA3 and output of NCS was demultiplexed and converted to FastQ format with Illumina *Bcl2fastq* v1.9.0.

For analysis, briefly, raw reads were trimmed with *cutadapt* (Martin, 2011) and *Trim Galore!* (http://www.bioinformatics.babraham.ac.uk/projects/trim_galore/) followed by quality control analysis via *FastQC* (Andrews, 2010) and *MultiQC* (Ewels, Magnusson, Lundin, & Kaller, 2016). Trimmed reads were aligned to hg38 assembly and indexed to GENCODE (v28) along with gene feature counting via *STAR* (Dobin et al., 2013). Low count genes with less than 1 count per sample on average were filtered prior to count normalization and differential gene expression (DGE) analysis by *DESeq2* with empirical Bayes shrinkage for fold-change estimation (Love, Anders, Kim, & Huber, 2015; Love, Huber, & Anders, 2014). Wald probabilities were corrected for multiple testing by independent hypothesis weighting (IHW) (Ignatiadis, Klaus, Zaugg, & Huber, 2016) for downstream analyses. In presented analyses, “log_2_FC” is the empirically observed log_2_ fold-change in expression between conditions, while “_s_log_2_FC” is a moderated log_2_ fold-change estimate that removes noise from low count genes using the *apeglm* shrinkage estimator as implemented in *DESeq2* (Zhu, Ibrahim, & Love, 2019). Pairwise comparisons between different DGE analyses and gene sets were initially filtered for genes with transcripts commonly detected in both cell populations.

### Histone peptide arrays

Anti-acetyl-H2A.Z (K4/K7) (D3V1I, Cell Signaling) antibody specificity was analyzed via histone peptide microarrays as previously described (Cornett, Dickson, & Rothbart, 2017) with minor modifications. Arrays were designed in *ArrayNinja* (Dickson, Cornett, Ramjan, & Rothbart, 2016) and printed using a 2470 Arrayer (Quanterix). All hybridization and wash steps were performed at ambient temperature. Slides were blocked with hybridization buffer (1X PBS [pH 7.6], 0.1% Tween, 5% BSA) for 30 minutes, then incubated with primary antibody diluted 1:1000 in hybridization buffer for 1 hour. Slides were washed 3X for 5 minutes with PBS, then probed with Alexa647-conjugated secondary antibody diluted 1:5000 in hybridization buffer for 30 minutes. Slides were washed 3X for 5 minutes with PBS, dipped in 0.1X PBS to remove salt, and spun dry. Slides were scanned on an InnoScan 1100 microarray scanner (Innopsys) and images were analyzed and quantified using *ArrayNinja*. Plots were generated in Prism (GraphPad). Each peptide antigen is printed six times per array, and each antibody was screened on two separate arrays.

### Chromatin immunoprecipitation (ChIP-seq) and analysis

Wild-type and lentiviral shRNA particle transduced 12Z cells were treated with 1% formaldehyde in growth media for 10 minutes at ambient temperature. Formaldehyde was quenched by the addition of 0.125 M Glycine and incubation for 5 minutes at room temperature, followed by PBS wash and scraping. 1*107 crosslinked cells were used for each ChIP, and each antibody and condition for ChIP was performed in duplicate. Chromatin from crosslinked cells was fractionated by digestion with micrococcal nuclease using the SimpleChIP Enzymatic Chromatin IP Kit (Cell Signaling) according to the manufacturer protocol, followed by 30 seconds of sonication. ChIP was then performed according to the SimpleChIP Enzymatic Chromatin IP Kit (Cell Signaling) with the addition of 5 mM sodium butyrate to preserve histone acetylation. To each 1.25 mL IP, the following antibodies were used: 1:125 anti-histone H3.3 (2D7-H1, abnova); 1:50 anti-histone H2A.Z-acetyl (K4/K7) (D3V1I, Cell Signaling); 1:250 anti-histone H2A.Z (ab4174, abcam); 1:50 anti-acetyl-histone H4 (06-866, Millipore); 1:125 anti-histone H4K16ac (Active Motif); 1:50 anti-CHD4 (D4B7, Cell Signaling); 1:250 anti-ZMYND8 (A302-089, Bethyl). Crosslinks were reversed with 0.4 mg/mL Proteinase K (ThermoFisher) and 0.2 M NaCl at 65 °C for 2 hours. DNA was purified using the ChIP DNA Clean & Concentrator Kit (Zymo).

Libraries for input and IP samples were prepared by the Van Andel Research Institute Genomics Core. 10 ng of material was used for input samples, and the entire precipitated sample was used for IPs. Libraries were generated using the KAPA Hyper Prep Kit (v5.16) (Kapa Biosystems). Prior to PCR amplification, end-repaired and A-tailed DNA fragments were ligated to IDT for Illumina UDI Adapters (IDT DNA Inc.). Quality and quantity of the finished libraries were assessed using a combination of Agilent DNA High Sensitivity chip (Agilent Technologies), QuantiFluor® dsDNA System (Promega) and Kapa Illumina Library Quantification qPCR assays (Kapa Biosystems). Individually indexed libraries were pooled, and 50 bp, paired-end sequencing (for ZMYND8, H3.3, H2A.Zac, and H4K16ac) or 100 bp, single-end sequencing (for CHD4, H2A.Z, and pan-H4ac) was performed on an Illumina NovaSeq 6000 sequencer using a 100-cycle sequencing kit (Illumina). Each library was sequenced to minimum read depth of 80 million reads per input library and 40 million reads per IP library. Base calling was performed by Illumina NCS v2.0, and NCS output was demultiplexed and converted to FastQ format with Illumina *Bcl2fastq* v1.9.0.

New and re-analyzed (differential) ChIP-seq experiments were analyzed as previously described (Wilson et al., 2020). Briefly, wild-type CHD4 and differential H2A.Z and pan-H4ac ChIP-seq experiments were analyzed as single-end libraries, while wild-type ZMYND8 and differential H3.3, H2A.Zac, and H4K16ac ChIP-seq were analyzed as paired-end libraries. Raw reads for IPs and inputs were trimmed with *cutadapt* (Martin, 2011) and *Trim Galore!* (http://www.bioinformatics.babraham.ac.uk/projects/trim_galore/) followed by quality control analysis via *FastQC* (Andrews, 2010) and *MultiQC* (Ewels et al., 2016). Trimmed reads were aligned to GRCh38.p12 reference genome (Schneider et al., 2017) via *Bowtie2* (Langmead & Salzberg, 2012) with flag ‘--very-sensitivè. Aligned reads were sorted and indexed with samtools (H. Li et al., 2009). Only properly-paired read fragments were retained for paired-end libraries via *samtools view* with flag ‘-f 3’ followed by sorting and indexing. For libraries intended for differential analyses, molecular complexity was then estimated from duplicate rates by *ATACseqQC* (Ou et al., 2018) and *preseqR* (Daley & Smith, 2013), and libraries were subsampled to equivalent molecular complexity within an experimental design based on these estimates with samtools. *Picard MarkDuplicates* (http://broadinstitute.github.io/picard/) was used to remove PCR duplicates, followed by sorting and indexing. *MACS2* (Y. Zhang et al., 2008) was used to call peaks on each ChIP replicate against the respective input control. For CHD4 and ZMYND8 IPs, *MACS2* called broadPeaks with FDR < 0.05 threshold and otherwise default settings. For H2A.Z and H2A.Zac IPs, *MACS2* called narrowPeaks with FDR < 0.05 threshold and flags ‘--nomodel --extsize 146’ to bypass model building. For H3.3, pan-H4ac, and H4K16ac IPs, MACS2 called broadPeaks with FDR < 0.05 threshold and flags ‘--nomodel --extsize 146’ to bypass model building. The resulting peaks were repeat-masked by ENCODE blacklist filtering and filtered for non-standard contigs (Amemiya, Kundaje, & Boyle, 2019). A naive overlapping peak set, as defined by ENCODE (Landt et al., 2012), was constructed by calling peaks on pooled replicates followed by *bedtools intersect* (Quinlan & Hall, 2010) to select for peaks of at least 50% overlap with each biological replicate.

ChIP-seq differential histone abundance analysis was performed with *csaw* (Lun & Smyth, 2016). First, a consensus peak set was constructed for each differential experiment from the union of replicate-intersecting, filtered *MACS2* peak regions called in each condition. ChIP reads were counted in these query regions by csaw, then filtered for low abundance peaks with average log_2_CPM > -3. When comparing ChIP libraries, any global differences in IP efficiency observed between the two conditions were considered a result of technical bias to ensure a highly conservative analysis. As such, we employed a loess-based local normalization to the peak count matrix, as is implemented in *csaw* (Lun & Smyth, 2016), to assume a symmetrical MA distribution. A design matrix was then constructed from one ‘condition’ variable, without an intercept term. The count matrix and loess offsets were then supplied to *edgeR* (M. D. Robinson, McCarthy, & Smyth, 2010) for estimating dispersions and fitting quasi-likelihood generalized linear models for differential abundance hypothesis testing. Nearby query regions were then merged up to 500 bp apart for a maximum merged region width of 5 kb, and the most significant probability was used to represent the merged region. Finally, FDR < 0.05 threshold was used to define significant differentially abundant regions.

### Chromatin state modeling and optimization

The same genome-wide chromatin 18-state map of 12Z cells with or without ARID1A depletion, constructed with *ChromHMM* (Ernst and Kellis 2012, 2017) using total RNA, ATAC, H3K4me1, H3K4me3, H3K18ac, H3K27ac, and H3K27me3 data (Wilson et al. 2020), was re-analyzed in Figure 2 and Figure 3 studies. A refined *ChromHMM* model was constructed with further addition of H3.3, H2A.Z, H2A.Zac (K4/K7), pan-H4ac (K5/K8/K12/K16), and H4K16ac features with some procedural modifications. In order to reduce technical confounders in differential chromatin state analysis between control and ARID1A-depleted cell types, we adopted an equalized binarization framework described by Fiziev et al. (Fiziev et al. 2017). Briefly, the *ChromHMM* chromosomal signal intermediate files during BAM binarization were saved and imported into R. Feature signal values were then background-subtracted by respective control signals when available (e.g. input chromatin for ChIP; does not occur for ATAC). For each feature and cell type, those (background-subtracted) signal values were ranked, and the top *n* ranked binarization calls are selected, where *n* is the lower number of calls among the two cell types for the given feature. The result is a new equalized binarization, where each feature has the same number of ‘present’ region calls in both cell types, per chromosome. As an example, if H3K18ac called 27,000 present regions on chromosome 1 in control cells and 35,000 present regions in ARID1A-depleted cells, then the top 27,000 regions are retained in both cell types. Chromatin state models from 5 to 40 states were then computed using the “concatenated” approach to unify both cell types for differential state comparisons. The new chromatin-state model was optimized at 25 states through a strategy devised by Gorkin et al. (Gorkin et al., 2020), which utilizes the *ChromHMM CompareModels* function to compare feature emission parameters from the 40-state (most complex) model against all other simpler models, as well as a k-means clustering of emission probabilities from all models together and analyzing the goodness of fit. See Figure 7—figure supplement 3 for related analyses. Across both strategies, 25 states was observed as a threshold for >95% median maximal state correlation and goodness of fit (between-cluster vs. total sum-of-squares) relative to the most complex model.

### Bioinformatics and statistics

The human endometrioma vs. control endometrium genome-wide expression (Illumina BeadChips) data set (Hawkins et al., 2011) was retrieved from GEO accession GSE23339 and analyzed via *GEO2R* and *limma* (Barrett et al., 2013; Davis & Meltzer, 2007; Ritchie et al., 2015). *biomaRt* was used for all gene nomenclature and mouse-human ortholog conversions (Smedley et al., 2009). The cumulative hypergeometric distribution was calculated in R for enrichment tests. *HOMER* was used to quantify sequencing reads across sets of genomic regions including heatmaps (Heinz et al., 2010). *GenomicRanges* functions were used to intersect and manipulate genomic coordinates (Lawrence et al., 2013). *IGV* (J. T. Robinson et al., 2011) was used for visualizing epigenomic data across hg38 loci as *MACS2* enrichment log-likelihood (logLR) for ChIP-seq and ATAC-seq or FPKM for RNA-seq. Hierarchical clustering by Euclidean distance and heatmaps were generated by *ComplexHeatmap* (Gu, Eils, & Schlesner, 2016). *ggplot2* was used for some plots in this study (Wickham, 2016). The statistical language R was used for various computing functions throughout this study (R Core Team, 2018).

## Authors’ Contributions

JJR and RLC conceptualized the study. JJR generated, analyzed, and curated all new experimental genomic data, led integrative computational and statistical analysis, and performed bioinformatic investigation of public data resources. MRW performed the cell cycle analysis. BA and CP assisted with experiments. MA led the library construction and sequencing efforts. JH and SBR performed histone peptide array experiments for antibody characterization. RLC supervised the study and provided funding. All authors read, critiqued, and approved the final manuscript.

## Acknowledgments

We thank the Van Andel Research Institute Genomics Core for providing sequencing facilities and services. We thank Dr. Jeremy Prokop for the use of his ÄKTA start FPLC to prepare linear glycerol gradients.

## Source Data Files

**Figure 3-figure supplement 1-source data 1.tif.** ARID1A Western Blot raw image as in Figure 3—figure supplement 1A and Figure 3—figure supplement 2.

**Figure 3-figure supplement 1-source data 2.tif.** β-actin Western Blot raw image as in Figure 3—figure supplement 1A and Figure 3—figure supplement 2.

**Figure 3-figure supplement 1-source data 3.tif.** ARID1A Western Blot raw image as in Figure 3—figure supplement 1B and Figure 3—figure supplement 2.

**Figure 3-figure supplement 1-source data 4.tif.** β-actin Western Blot raw image as in Figure 3—figure supplement 1B and Figure 3—figure supplement 2.

**Figure 3-figure supplement 1-source data 5.tif.** Histone H3.3 Western Blot raw image as in Figure 3—figure supplement 1B and Figure 3—figure supplement 2.

**Figure 3-figure supplement 1-source data 6.tif.** Total histone H3 Western Blot raw image (right blot) as in Figure 3—figure supplement 1B and Figure 3—figure supplement 2.

**Figure 4-source data 1.tif.** Grayscale, high contrast histone H3.3 (left) and total histone H3 (right) Western Blot raw image as in Figure 4B and Figure 4—figure supplement 2. Presented image is flipped vertically.

**Figure 4-source data 2.tif.** Grayscale, low contrast histone H3.3 (left) and total histone H3 (right) Western Blot raw image as in Figure 4B and Figure 4—figure supplement 2. Presented image is flipped vertically.

**Figure 4-source data 3.tif.** Dual-color fluorescent histone H3.3 (left) and total histone H3 (right) Western Blot raw image as in Figure 4—figure supplement 2. Presented image is flipped vertically.

**Figure 5-source data 1.tif.** ARID1A Western Blot raw image as in Figure 5C and Figure 5— figure supplement 1.

**Figure 5-source data 2.tif.** CHD4 Western Blot raw image as in Figure 5C and Figure 5—figure supplement 1.

**Figure 5-source data 3.tif.** ZMYND8 Western Blot raw image as in Figure 5C and Figure 5— figure supplement 1.

**Figure 5-source data 4.tif.** CHD4 Western Blot raw image as in Figure 5D and Figure 5— figure supplement 1.

**Figure 5-source data 5.tif.** ARID1A Western Blot raw image as in Figure 5D and Figure 5— figure supplement 1.

**Figure 5-source data 6.tif.** ZMYND8 Western Blot raw image as in Figure 5D and Figure 5— figure supplement 1.

**Figure 5-source data 7.tif.** ARID1A Western Blot raw image as in Figure 5E and Figure 5— figure supplement 1.

**Figure 5-source data 8.tif.** BRG1 Western Blot raw image as in Figure 5E and Figure 5—figure supplement 1.

**Figure 5-source data 9.tif.** BAF155 Western Blot raw image as in Figure 5E and Figure 5— figure supplement 1.

**Figure 5-source data 10.tif.** ZMYND8 Western Blot raw image as in Figure 5E and Figure 5— figure supplement 1.

**Figure 5-source data 11.tif.** CHD4 Western Blot raw image as in Figure 5E and Figure 5— figure supplement 1.

**Figure 5-source data 12.tif.** HDAC1 Western Blot raw image as in Figure 5E and Figure 5— figure supplement 1.

**Figure 7-figure supplement 6-source data 1.tif.** ZMYND8 Western Blot raw image as in Figure 7—figure supplement 6A and Figure 7—figure supplement 9.

**Figure 7-figure supplement 6-source data 2.tif.** β-actin Western Blot raw image as in Figure 7—figure supplement 6A and Figure 7—figure supplement 9.

**Figure 7-figure supplement 6-source data 3.tif.** CHD4 Western Blot raw image as in Figure 7—figure supplement 6D and Figure 7—figure supplement 9.

**Figure 7-figure supplement 6-source data 4.tif.** β-actin Western Blot raw image as in Figure 7—figure supplement 6D and Figure 7—figure supplement 9.

---

**Figure 1—figure supplement 1.**
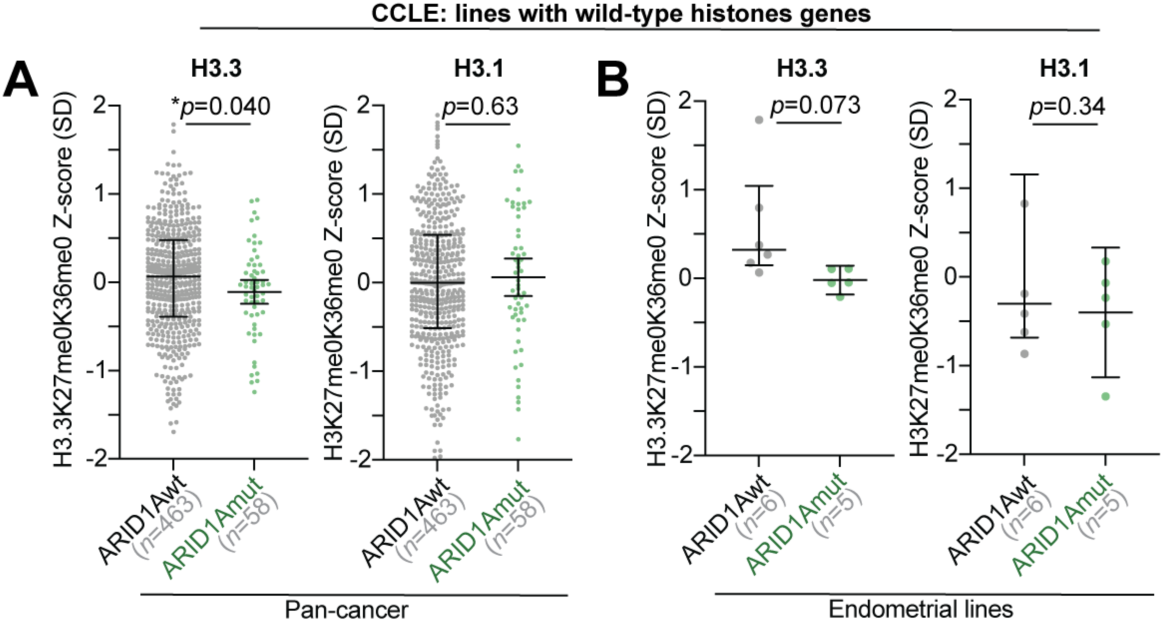
H3.3 analysis in histone wild-type lines. Repeated analysis of H3.3 and H3.1 abundance in CCLE lines that are wild-type for all 74 human histone genes annotated by Nacev et al. Statistic is two-tailed, unpaired Welch’s *t*-test. **A**, H3.3 (left) and H3.1 (right) peptide abundances in ARID1A mutant vs. wild-type pan-cancer lines, as in Figure 1C. **B**, H3.3 (left) and H3.1 (right) peptide abundances in ARID1A mutant vs. wild-type endometrial cancer lines, as in Figure 1D.

**Figure 3—figure supplement 1.**
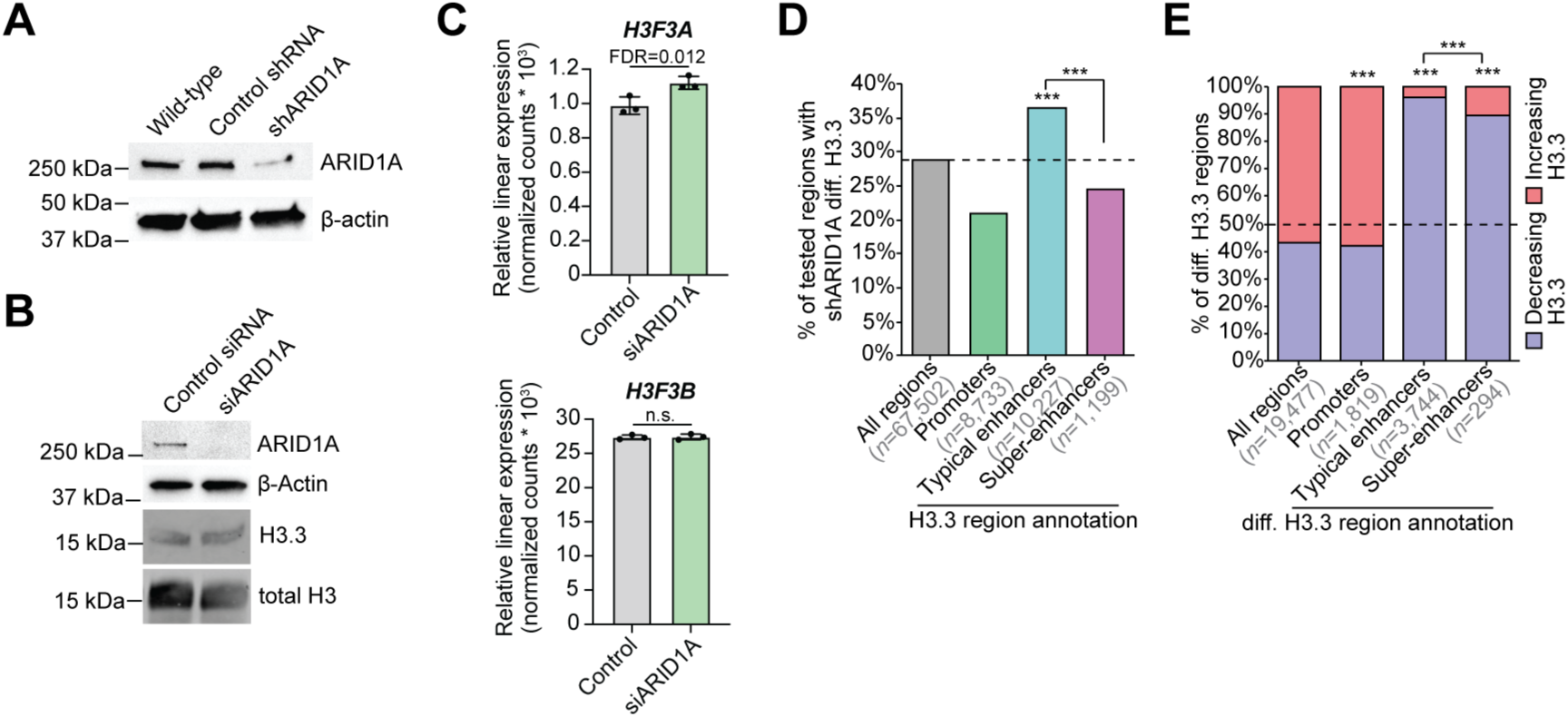
Supplemental ARID1A knockdown differential H3.3 data. **A**, ARID1A immunoblot, compared to β-actin loading control, in 12Z cells treated with lentiviral shRNA particles to acutely knockdown ARID1A compared to non-targeting control shRNA. Wild-type condition shown as reference. **B**, Immunoblot for H3.3 and ARID1A, compared to total H3 and β-actin loading controls, respectively, in 12Z cells treated with siRNA to acutely knockdown ARID1A (siARID1A). **C**, RNA-seq linear gene expression data (normalized counts) for H3.3-encoding gene isoforms, *H3F3A* and *H3F3B*, following ARID1A knockdown by siRNA (siARID1A). **D**, Enrichment of regions displaying shARID1A significant differential H3.3 abundance among promoters, typical enhancers, and super-enhancers compared to all tested H3.3 regions. Gene promoters are defined as within 3 kb of a TSS. Enhancers are defined as ATAC+ H3K27ac peaks located >3 kb from a TSS. Super-enhancers were further distinguished from typical enhancers by *ROSE*. Statistics are hypergeometric enrichment and pairwise two-tailed Fisher’s exact test. **E**, Distribution of significantly increasing vs. decreasing genomic H3.3 with ARID1A knockdown among the classes described in **C**. Statistics are hypergeometric enrichment and pairwise two-tailed Fisher’s exact test.

**Figure 3—figure supplement 2.**
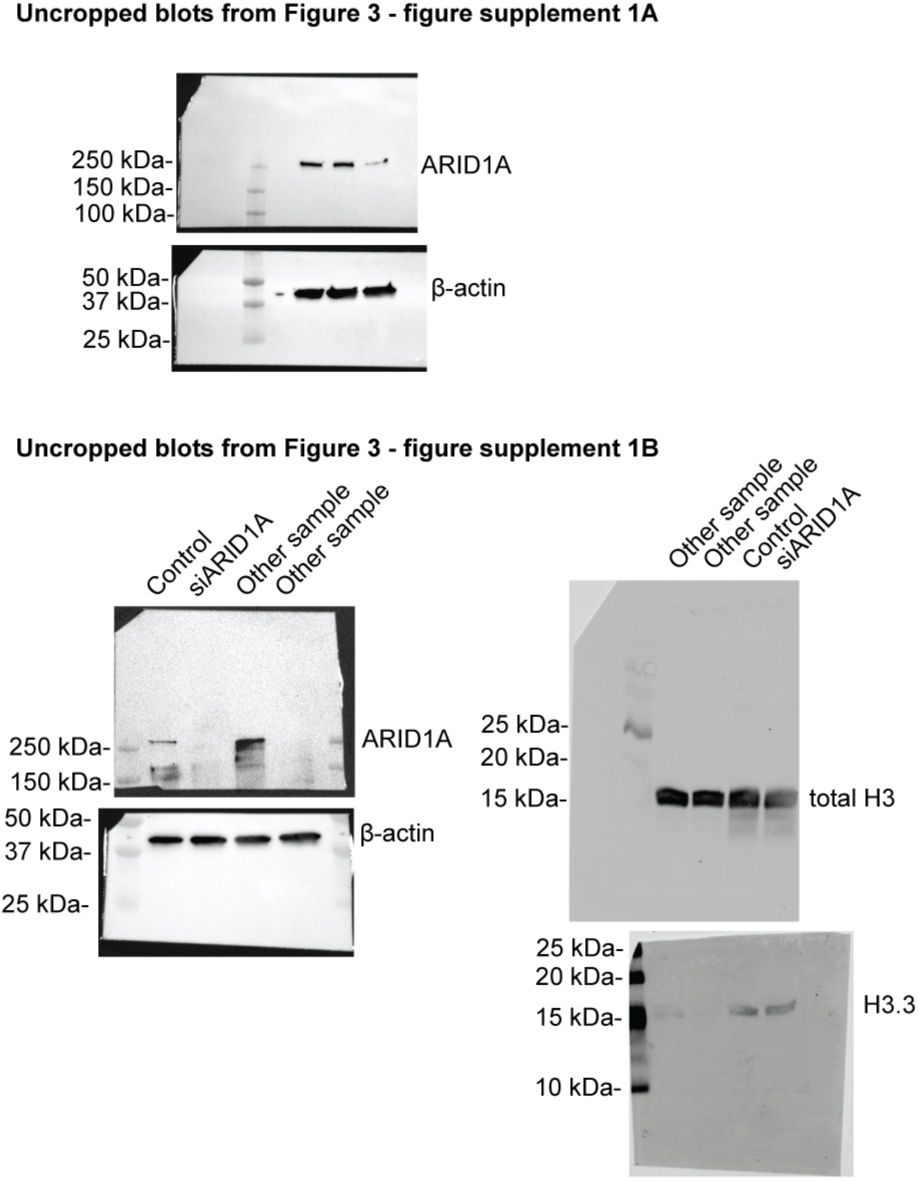
Uncropped Western blots from Figure 3—figure supplement 1.

**Figure 4—figure supplement 1.**
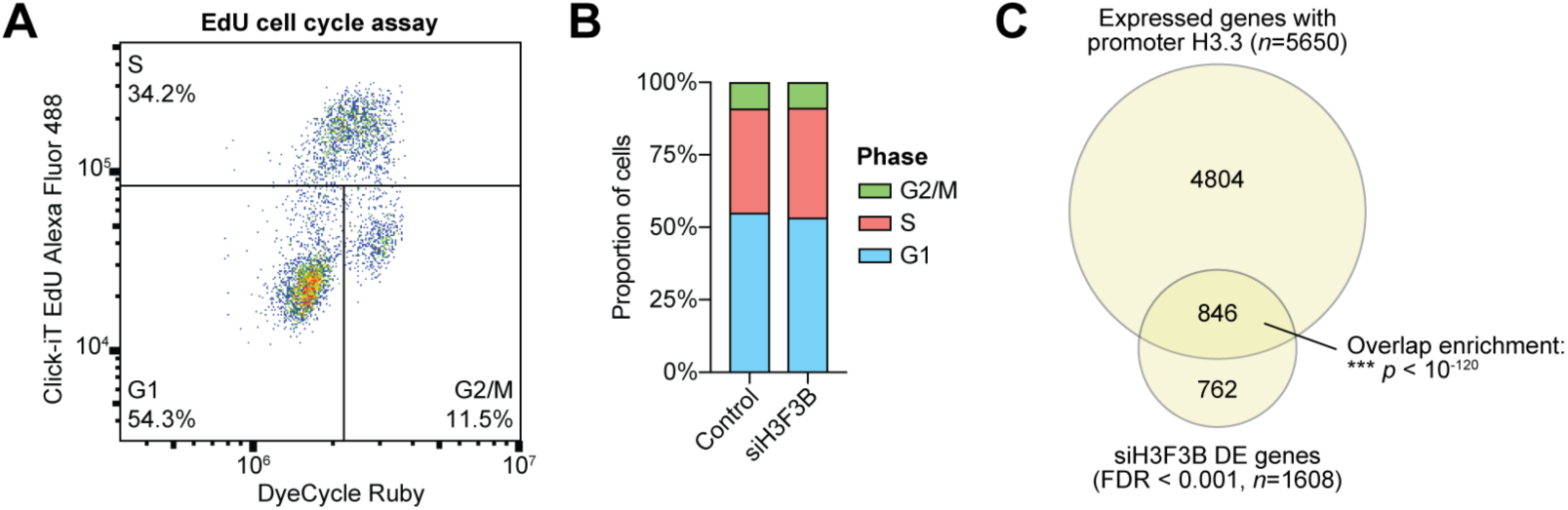
H3.3 knockdown functional analysis. **A**, Representative cell cycle analysis by EdU labeling and DyeCycle ruby staining following 72 hours siRNA trasnfection. **B**, Quantification of cell cycle phases (G1, S, and G2/M) in control non-targeting siRNA treated and siH3F3B (H3.3 knockdown) treated 12Z cells. **C**, Euler diagram displaying overlap of genes with detected promoter H3.3 by ChIP-seq (*MACS2*, FDR < 0.05) and genes with altered expression following H3.3 knockdown by RNA-seq (*DESeq2*, FDR < 0.001). Statistic is hypergeometric enrichment.

**Figure 4—figure supplement 2.**
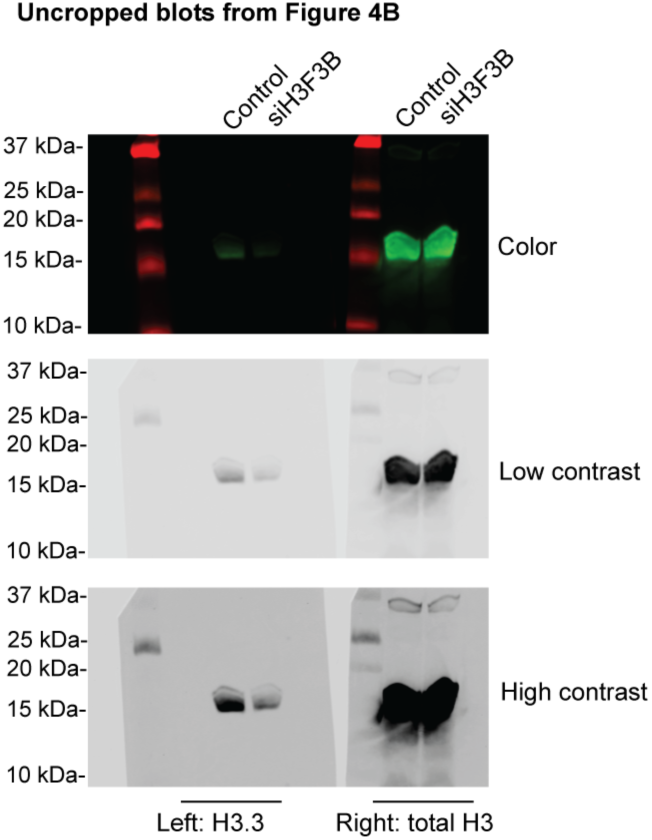
Uncropped Western blots from Figure 4. Color image is presented for molecular weight marker reference.

**Figure 5—figure supplement 1.**
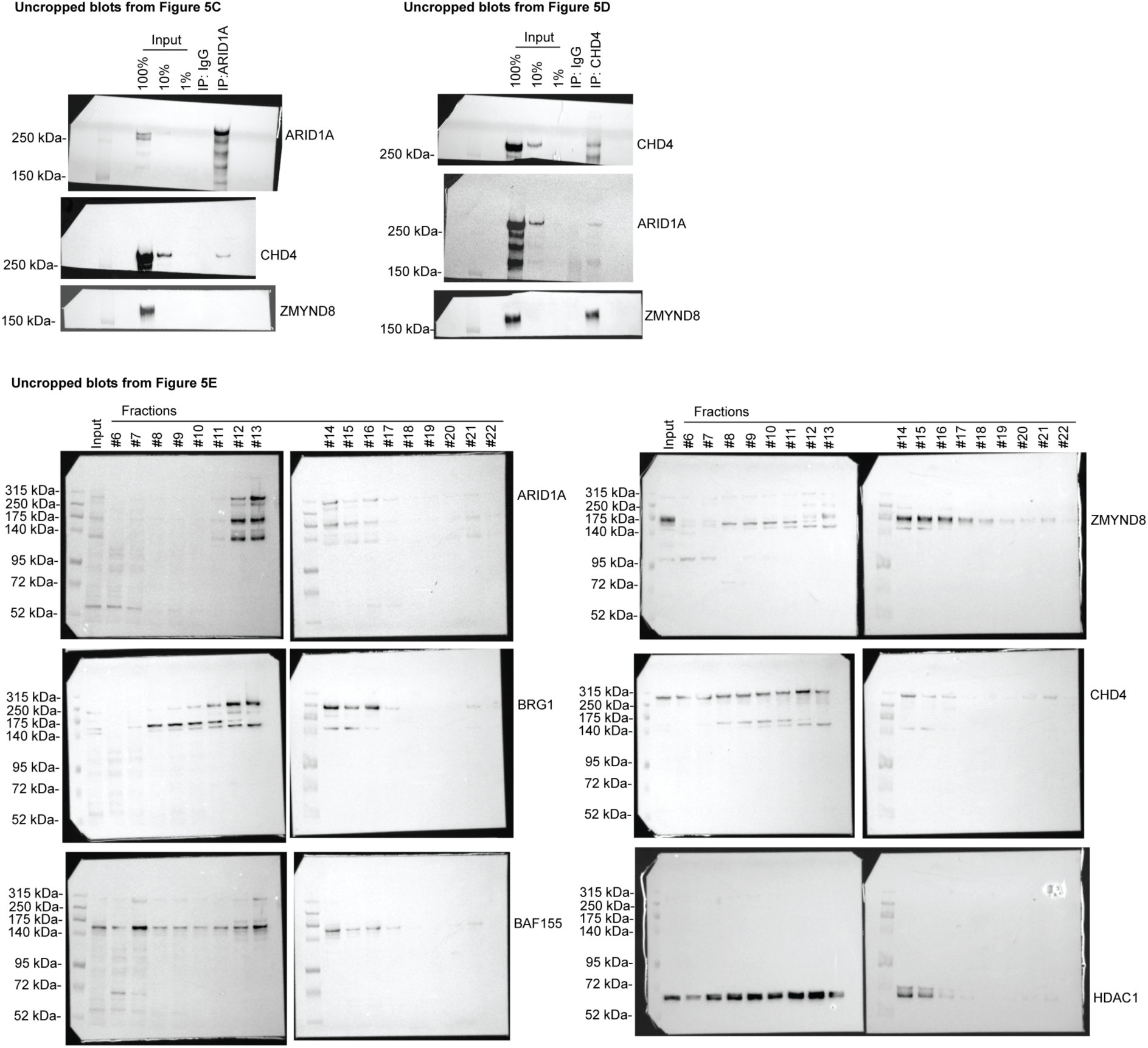
Uncropped Western blots from Figure 5.

**Figure 7—figure supplement 1.**
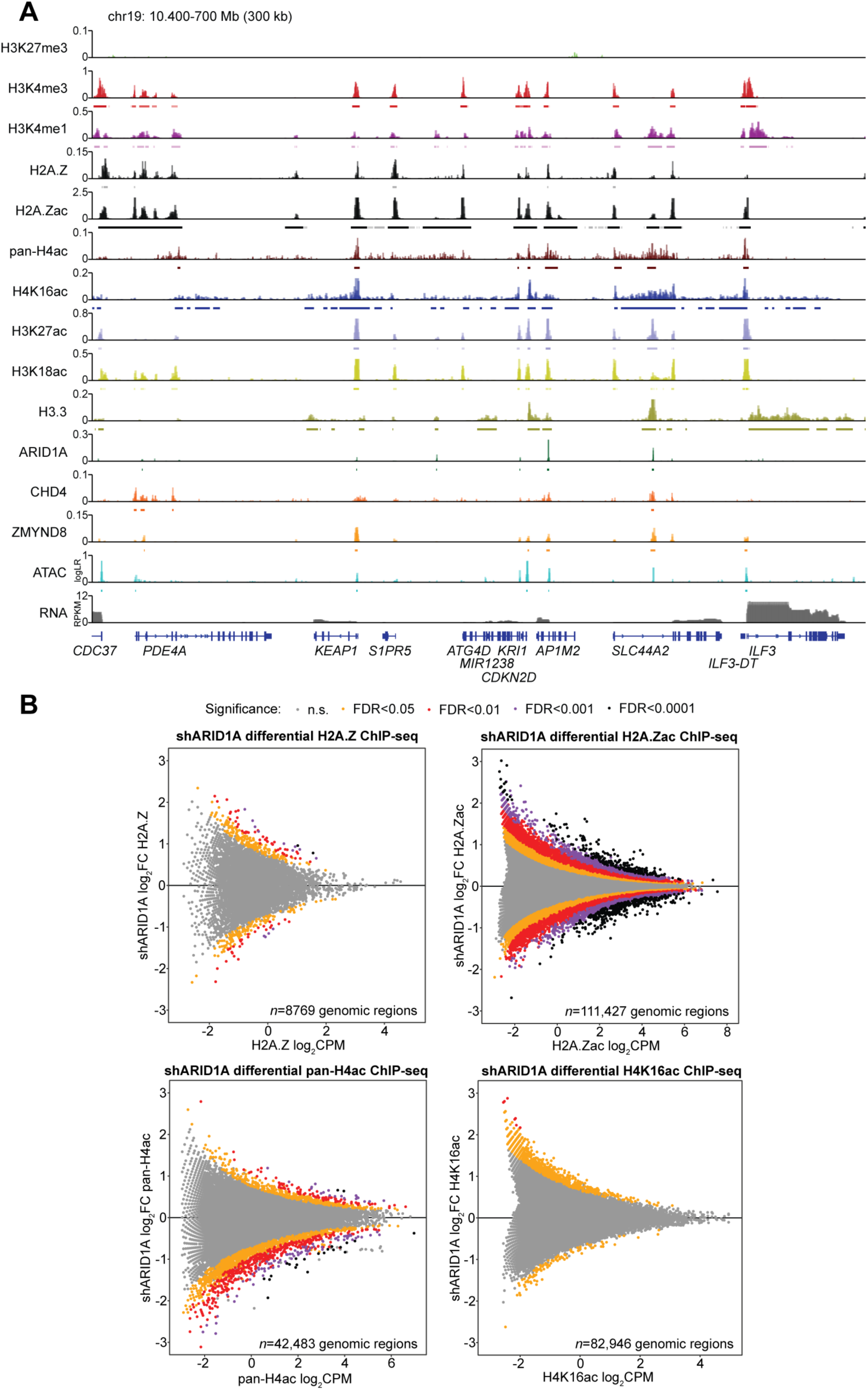
Additional chromatin features profiled in 12Z cells. **A**, Example hg38 locus on chromosome 19 depicting all analyzed chromatin features in this study. *y*-axis is log-likelihood ratio (logLR) of assay signal (compared to input chromatin for ChIP-seq or background genome for ATAC-seq) or RPKM for total RNA. Small bars under tracks indicate significant peak detection by *MACS2* (FDR < 0.05). **B**, MA plots for new shARID1A vs. control differential ChIP-seq experiments: H2A.Z, H2A.Zac (K4/K7), pan-H4ac (K5/K8/K12/K16), and H4K16ac. Genomic regions are colored by significance (*csaw*/*edgeR*).

**Figure 7—figure supplement 2.**
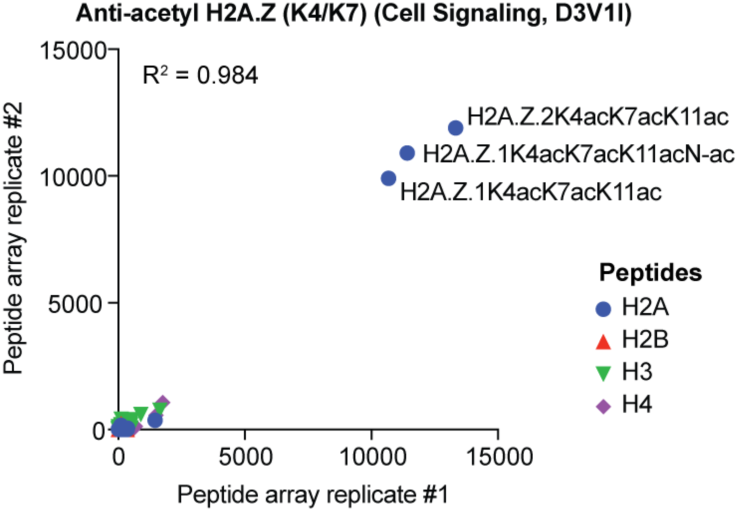
Peptide specificity of anti-acetyl-H2A.Z (K4/K7). Hybridizaiton results for two independent histone peptide microarray replicates probed with anti-acetyl-H2A.Z (H2A.Zac) (Cell Signaling, D3V1I), demonstrating clear specificity for acetylated H2A.Z peptides. The three dominantly detected H2A.Zac peptides are labeled.

**Figure 7—figure supplement 3.**
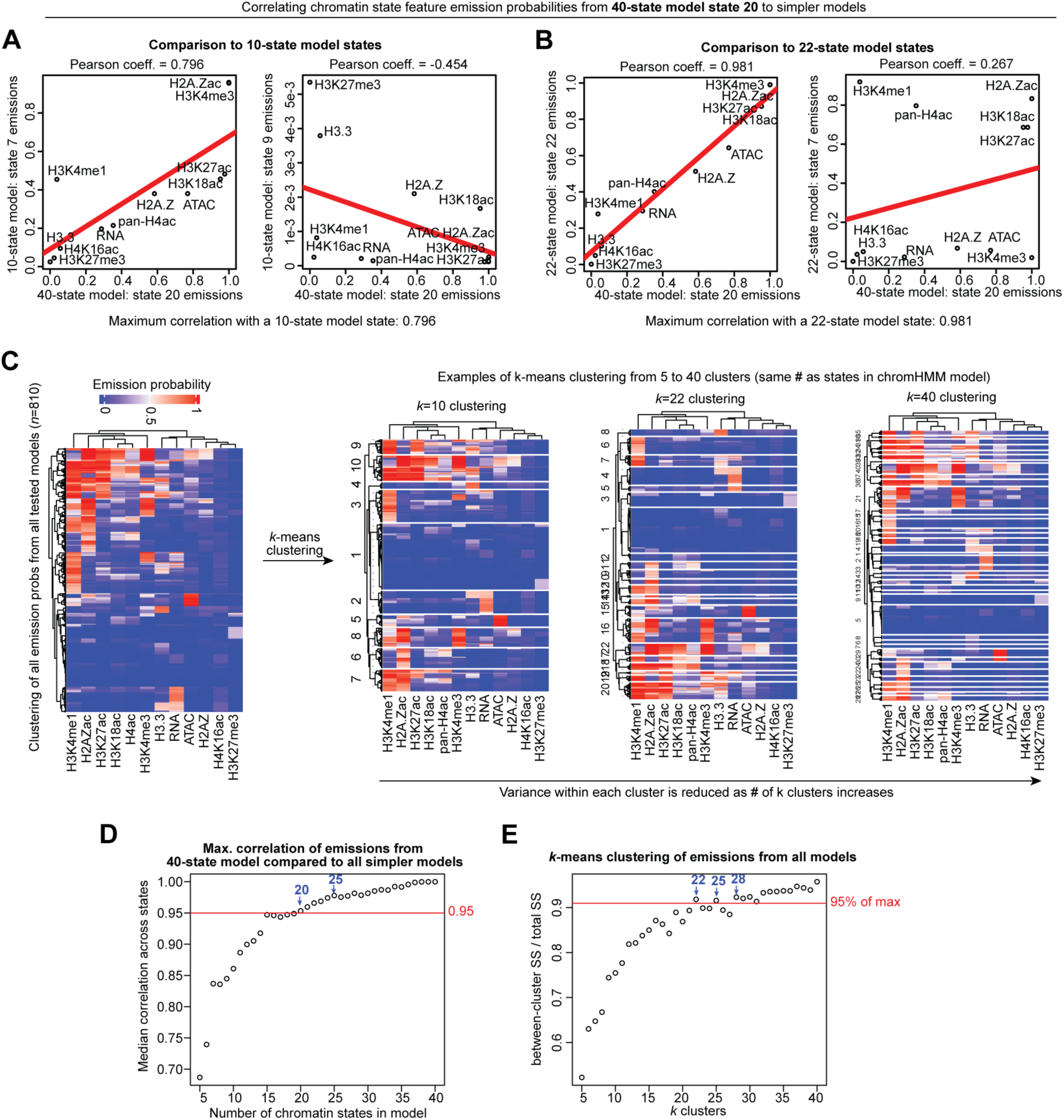
ChromHMM model optimization. Examples and results of quantitative strategies used to select optimal ChromHMM model as devised by Gorkin et al. See Materials and Methods for further details and procedural description. **A**-**B**, Correlating chromatin state feature emission probabilities from the most complex model (40-state) to all simpler models. **A**, Comparison to 10-state model states. The maximum Pearson correlation coefficient of 40-state model state 20 with a 10-state model state is 0.796. **B**, Comparison to 22-state model states. The maximum Pearson correlation coefficient of 40-state model state 20 with a 22-state model state is 0.981. **C**, Clustered heatmaps of all emission probabilities from all tested models (left*, n* = 810 total modeled chromatin states), followed by examples of *k*-means clustering (right), *k*-means clustering was performed from *k* = 5 to *k* = 40 clusters (the same number of states in the tested chromatin state models) for measurements of goodness of fit by sum-of-squares (between-cluster vs. total). Variance within each cluster is reduced as the number of *k* clusters increases. **D**, First strategy model selection plot (as in **A**-**B**) based on the maximum correlation of emissions from the most complex model (40-state) compared to all simpler models. The *x*-axis is the simpler model used for comparison, and the *y*-axis is the median of maximum correlation coefficients across all 40-state model states compared to states within the simpler model. 95% threshold is represented by the red line. **E**, Second strategy model selection plot (as in **C**) based on the goodness of fit for all simpler models relative to the most complex model (40-state). The *y*-axis is the amount of variance explained by clustering, i.e. the ratio of between-cluster sum-of-squares to total sum-of-squares, relative to *k* = 40 clusters (the number of chromatin states in the most complex model). 95% threshold is represented by the red line.

**Figure 7—figure supplement 4.**
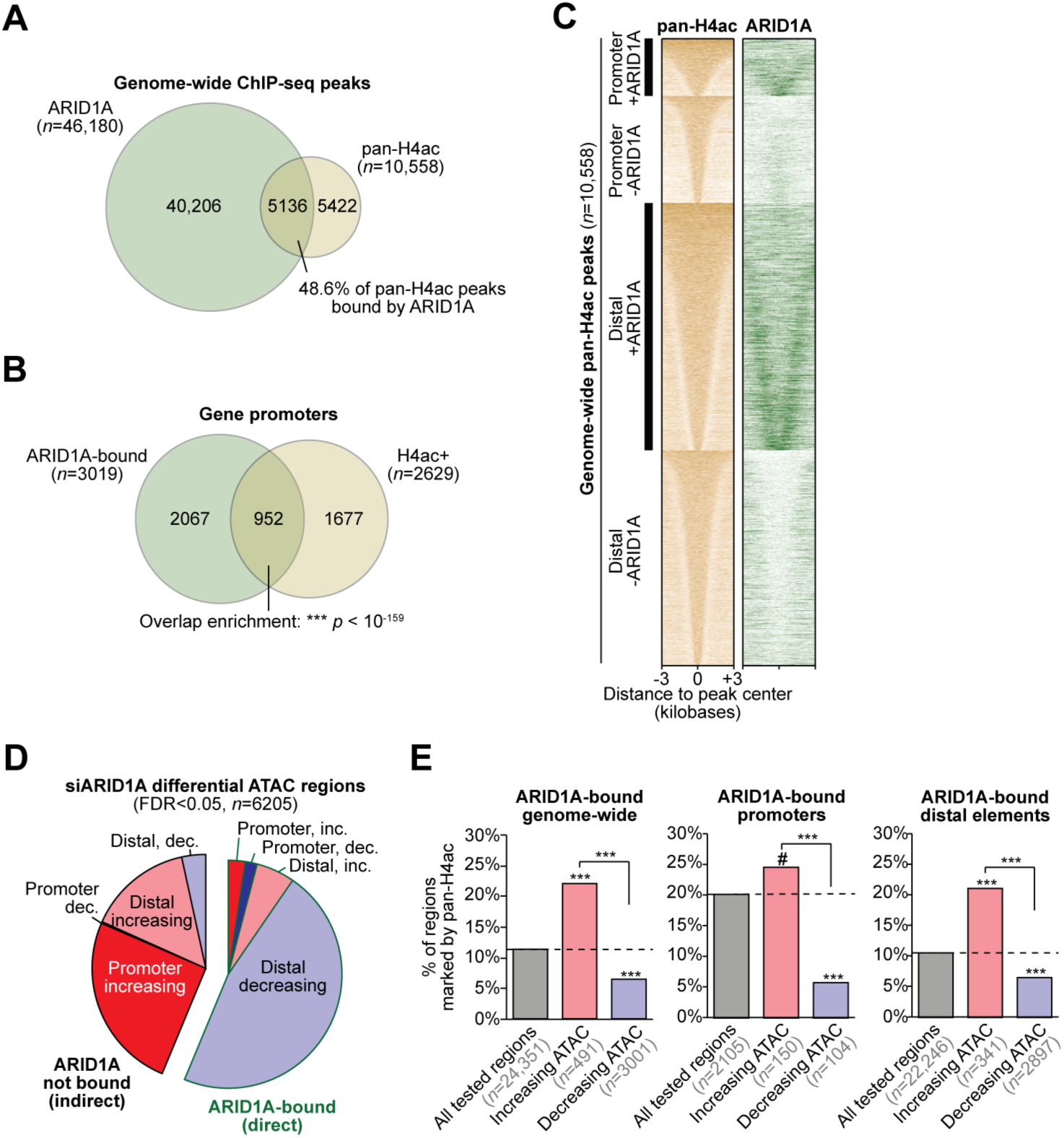
Chromatin accessibility repressed by ARID1A is associated with H4 acetylation. **A**, Euler diagram displaying overlapping genome-wide ChIP-seq peaks (called by *MACS2*) for pan-H4ac compared to ARID1A. 48.6% of pan-H4ac peaks are co-marked by ARID1A binding. **B**, Euler diagram displaying overlapping gene promoters marked by pan-H4ac (H4ac+) and ARID1A binding. Statistic is hypergeometric enrichment test. **C**, Heatmap displaying ChIP-seq signal across 10,558 genome-wide pan-H4ac ChIP-seq peaks. Signal is quantified as ChIP – Input. Peaks are ranked by overall pan-H4ac signal and stratified by ARID1A binding and promoter (<3 kb from a TSS) vs. distal (>3 kb from a TSS). **D**, Annotation and directional breakdown of 6205 significant (FDR < 0.05) differentially accessible genomic regions, measured by ATAC-seq, following ARID1A depletion via siRNA (siARID1A). Regions are further segregated based on ARID1A binding status. **E**, Association of pan-H4ac with siARID1A differential ATAC regions directly bound by ARID1A and separated by direction of accessibility change genome-wide (left), at promoters (center), and at distal elements (right). Statistic is hypergeometric enrichment test and two-tailed Fisher’s exact test. # *p* < 0.10, *** *p* < 0.001.

**Figure 7—figure supplement 5.**
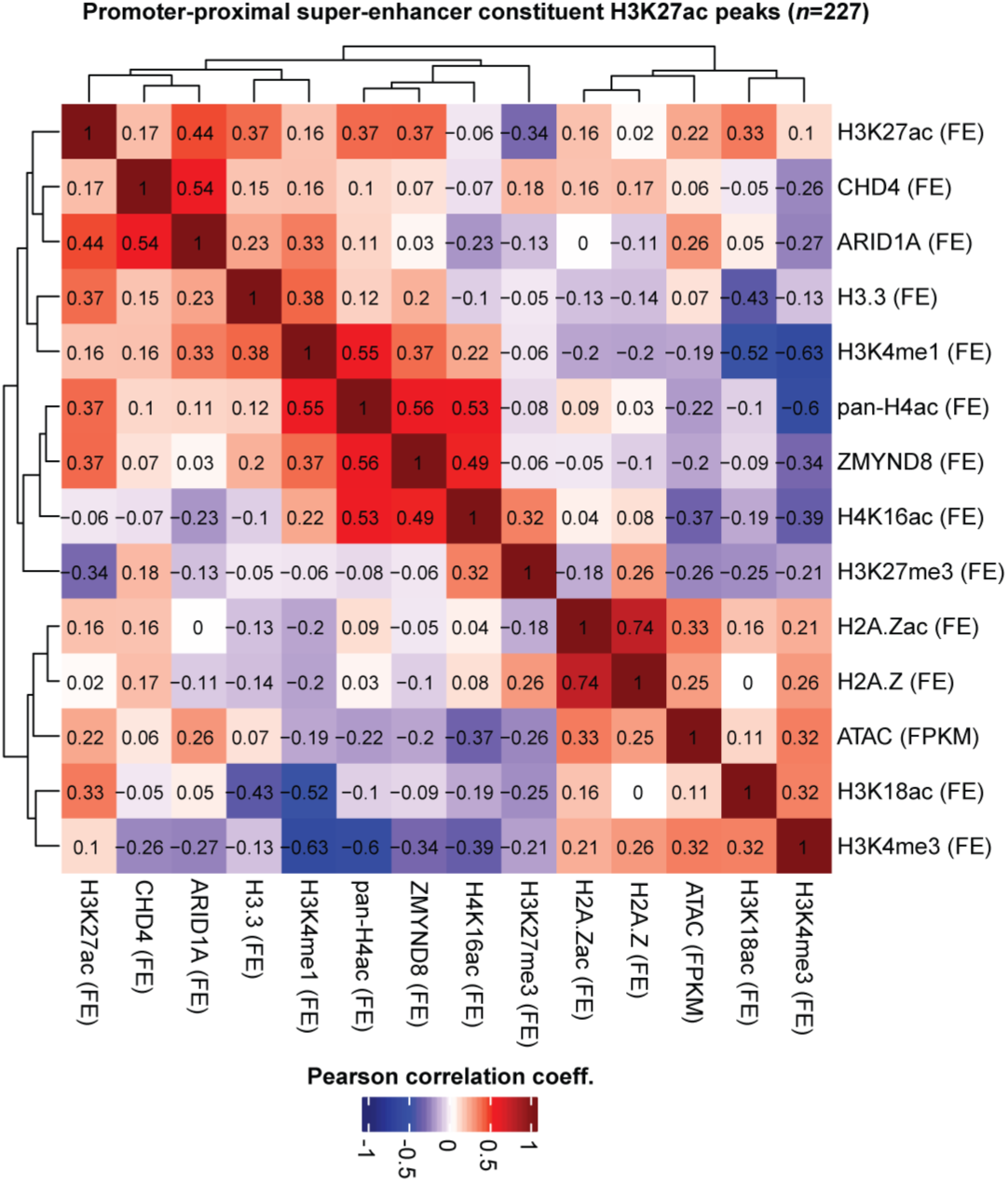
Chromatin feature correlation across promoter-proximal super-enhancers. Clustered heatmap of Pearson correlation coefficients for all measured chromatin features (aside from total RNA) across 227 promoter-proximal super-enhancer constituent H3K27ac peaks. These active super-enhancer-like regions (distinguished by *ROSE*) overlap the 3 kb promoter region flanking a gene TSS. ChIP-seq assays are quantified as ChIP/input chromatin fold-enrichment (FE), and ATAC-seq is quantified as FPKM.

**Figure 7—figure supplement 6.**
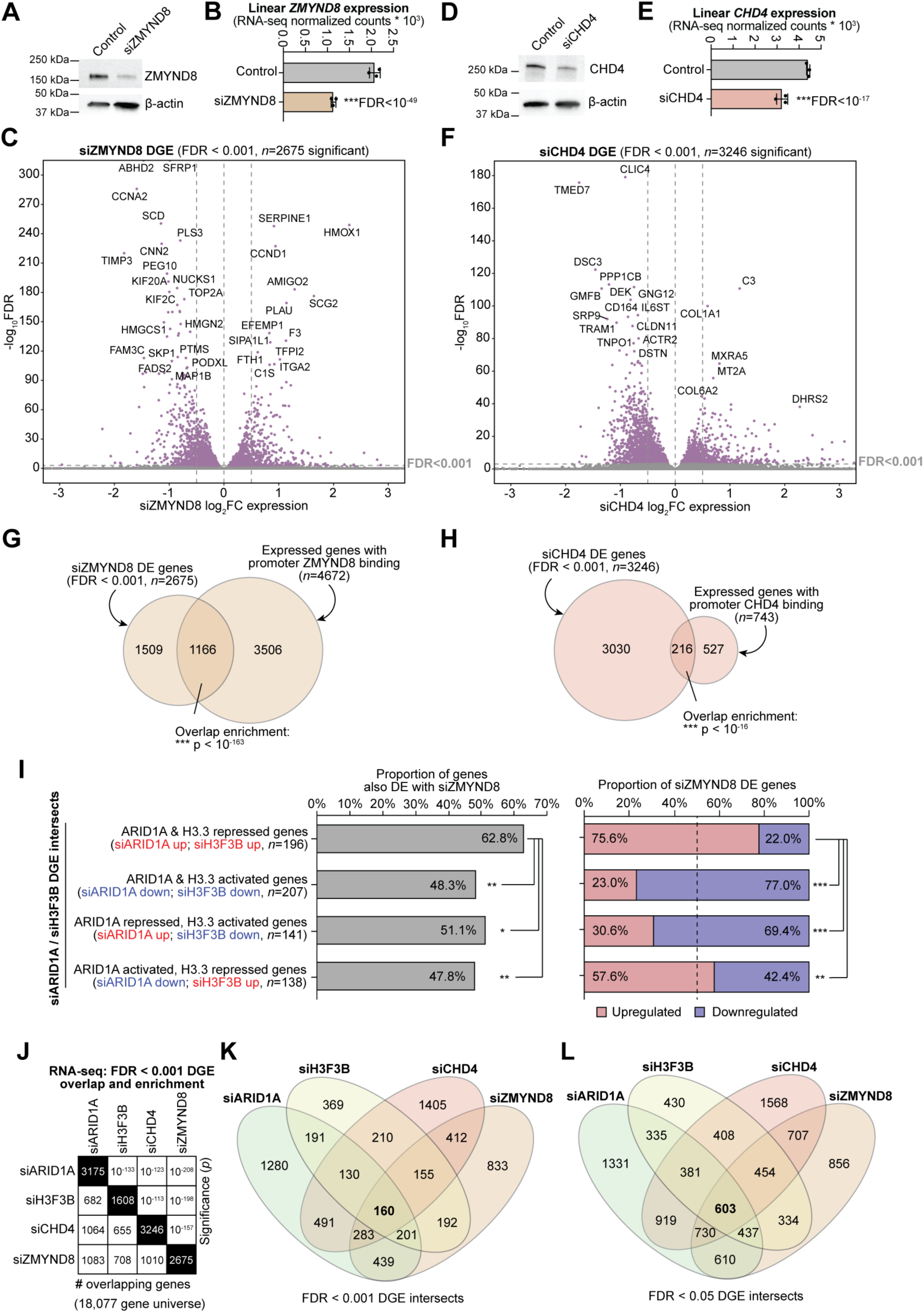
siCHD4/siZMYND8 functional analysis. **A**, Immunoblot for (top) ZMYND8 compared to (bottom) β-actin loading control in 12Z cells treated with non-targeting control siRNA or siZMYND8 (ZMYND8 knockdown). **B**, RNA-seq expression of ZMYND8 in control and siZMYND8 cells. Statistic is FDR-adjusted *DESeq2* Wald test. **C**, Volcano plot for differential gene expression between siZMYND8 and control cells. FDR < 0.001 was used as a significance threshold. Top significant ZMYND8-dependent genes are labeled. **D**-**F**, CHD4 siRNA knockdown (siCHD4) framework and RNA-seq analysis as in **A**-**C**. **G**, Euler diagram displaying overlap between siZMYND8 differential gene expression (DGE) and ZMYND8 promoter-bound genes. Statistic is hypergeometric enrichment test. **H**, Euler diagram displaying overlap between siCHD4 DGE and CHD4 promoter-bound genes. Statistic is hypergeometric enrichment test. **I**, Left, association of siZMYND8 DGE among ARID1A-H3.3 co-regulated gene classes described in Fig. 3F-H. Right, distribution of significantly upregulated (ZMYND8 repressed) vs. downregulated (ZMYND8 activated) siZMYND8 DE genes among ARID1A-H3.3 co-regulated gene classes. Statistic is two-tailed Fisher’s exact test. **J**, RNA-seq DGE (FDR < 0.001) overlap and enrichment across the four analyzed knockdown conditions: siARID1A, siH3F3B, siCHD4, and siZMYND8. The black cell diagonal represents the number of total significant DE genes in that condition. The bottom-left triangle displays the number of overlapping DE genes between pairwise knockdowns. The upper-right triangle displays the overlap enrichment significance by hypergeometric enrichment test. A reduced 18,077 gene set universe was used that contains genes with detected expression in all analyzed conditions. **K**, Overlap of FDR < 0.001 DGE sets across the four analyzed knockdown conditions. **L**, Overlap of FDR < 0.05 DGE sets across the four analyzed knockdown conditions, corresponding with Fig. 6F.

**Figure 7—figure supplement 7.**
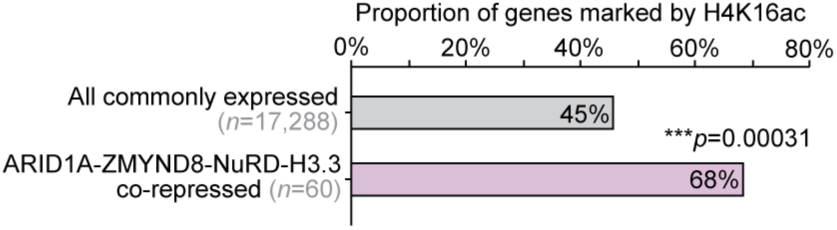
H4K16ac enrichment at repressed mechanistic genes. Enrichment of H4K16ac at gene promoters or gene bodies of ARID1A-H3.3-ZMYND8-CHD4 co-repressed genes—i.e. genes that are upregulated (FDR < 0.05) following treatment with siARID1A, siH3F3B, siZMYND8, and siCHD4—compared to all expressed genes with chromatin data. Statistic is hypergeometric enrichment test.

**Figure 7—figure supplement 8.**
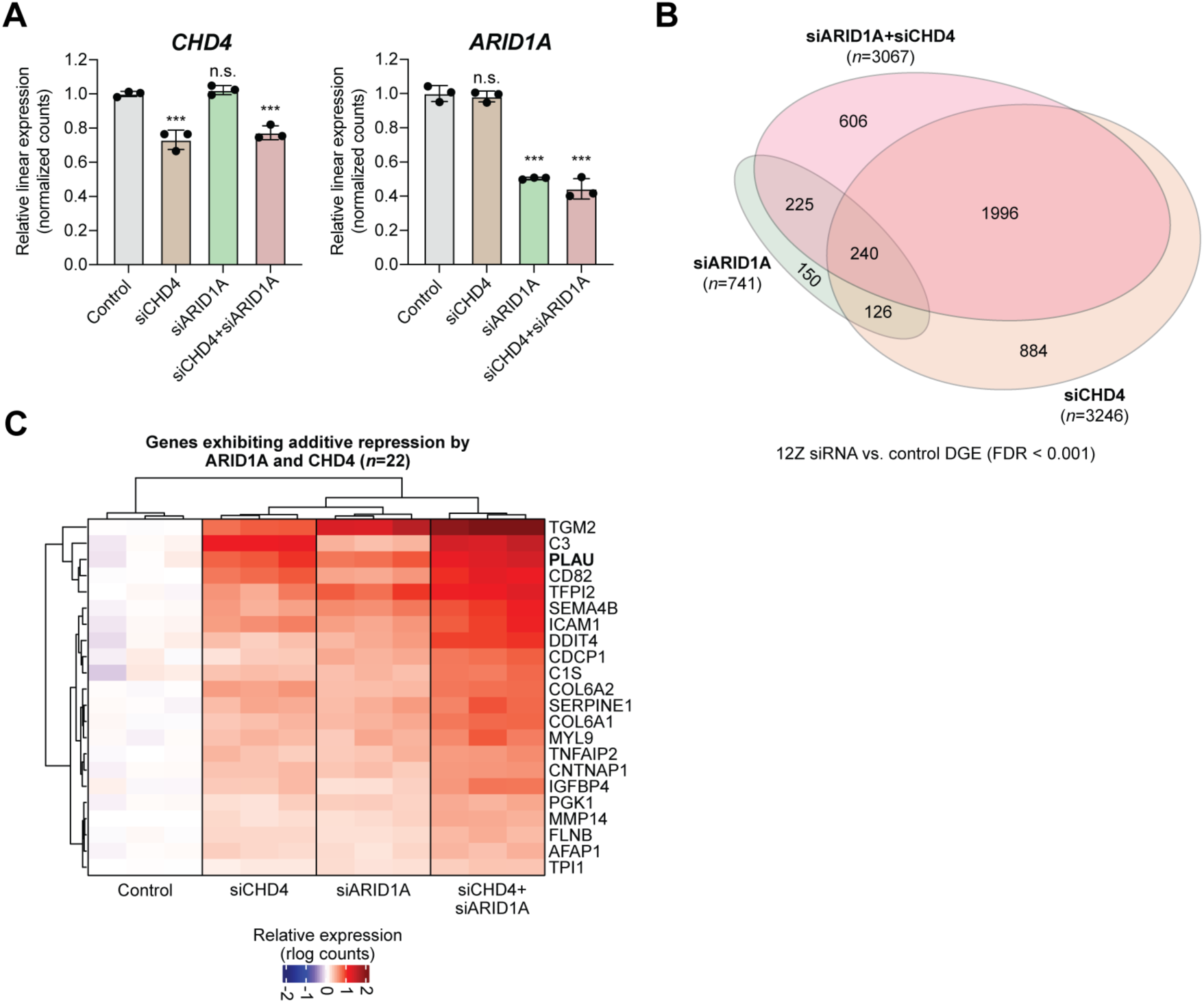
Additive transcriptional repression by ARID1A and CHD4. RNA-seq analysis of an ARID1A ± CHD4 co-knockdown experiment (*n* = 3 per condition). **A**, Relative linear RNA-seq expression for (left) CHD4 and (right) ARID1A among siRNA treated conditions. Statistic is *DESeq2* FDR-adjusted Wald test. **B**, Euler diagram displaying DGE (FDR < 0.001) overlap of siCHD4, siARID1A, and siARID1A+siCHD4 conditions compared to control cells. **C**, Clustered heatmap of expression alterations (rlog) at 22 genes displaying additive repression by ARID1A and CHD4. These genes are significantly upregulated in siCHD4, siARID1A, and siARID1A+siCHD4 conditions compared to control cells and siARID1A+siCHD4 vs. single-knockdown conditions. *** FDR < 0.001

**Figure 7—figure supplement 9.**
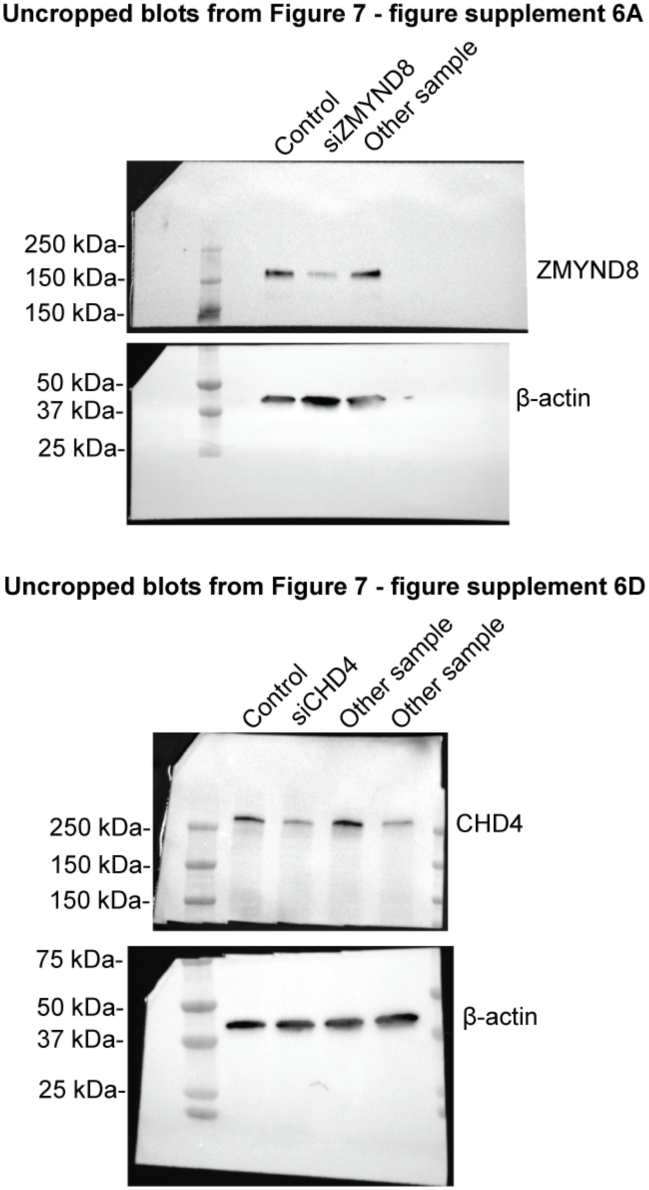
Uncropped Western blots from Figure 7—figure supplement 6.

## References

Adhikary, S., Sanyal, S., Basu, M., Sengupta, I., Sen, S., Srivastava, D. K., . . . Das, C. (2016). Selective Recognition of H3.1K36 Dimethylation/H4K16 Acetylation Facilitates the Regulation of All-trans-retinoic Acid (ATRA)-responsive Genes by Putative Chromatin Reader ZMYND8. Journal of Biological Chemistry, 291(6), 2664–2681. doi:10.1074/jbc.M115.679985

Ahmad, K., & Henikoff, S. (2002). The histone variant H3.3 marks active chromatin by replication-independent nucleosome assembly. Mol Cell, 9(6), 1191–1200. doi:10.1016/s1097-2765(02)00542-7

Alver, B. H., Kim, K. H., Lu, P., Wang, X., Manchester, H. E., Wang, W., . . . Roberts, C. W. (2017). The SWI/SNF chromatin remodelling complex is required for maintenance of lineage specific enhancers. Nat. Commun., 8, 14648. doi:10.1038/ncomms14648

Amemiya, H. M., Kundaje, A., & Boyle, A. P. (2019). The ENCODE Blacklist: Identification of Problematic Regions of the Genome. Sci Rep, 9(1), 9354. doi:10.1038/s41598-019-45839-z

Andrews, S. (2010). FastQC: A Quality Control Tool for High Throughput Sequence Data. Retrieved from http://www.bioinformatics.babraham.ac.uk/projects/fastqc

Anglesio, M. S., Papadopoulos, N., Ayhan, A., Nazeran, T. M., Noe, M., Horlings, H. M., . . . Shih, I. M. (2017). Cancer-Associated Mutations in Endometriosis without Cancer. New England Journal of Medicine, 376(19), 1835–1848. doi:10.1056/NEJMoa1614814

Barrett, T., Wilhite, S. E., Ledoux, P., Evangelista, C., Kim, I. F., Tomashevsky, M., . . . Soboleva, A. (2013). NCBI GEO: archive for functional genomics data sets--update. Nucleic Acids Res, 41(Database issue), D991–995. doi:10.1093/nar/gks1193

Barutcu, A. R., Lajoie, B. R., Fritz, A. J., McCord, R. P., Nickerson, J. A., van Wijnen, A. J., . . . Imbalzano, A. N. (2016). SMARCA4 regulates gene expression and higher-order chromatin structure in proliferating mammary epithelial cells. Genome Res, 26(9), 1188–1201. doi:10.1101/gr.201624.115

Basta, J., & Rauchman, M. (2015). The nucleosome remodeling and deacetylase complex in development and disease. Transl Res, 165(1), 36–47. doi:10.1016/j.trsl.2014.05.003

Bottardi, S., Mavoungou, L., Pak, H., Daou, S., Bourgoin, V., Lakehal, Y. A., . . . Milot, E. (2014). The IKAROS interaction with a complex including chromatin remodeling and transcription elongation activities is required for hematopoiesis. PLoS Genet, 10(12), e1004827. doi:10.1371/journal.pgen.1004827

Bui, C. B., Le, H. K., Vu, D. M., Truong, K. D., Nguyen, N. M., Ho, M. A. N., & Truong, D. Q. (2019). ARID1A-SIN3A drives retinoic acid-induced neuroblastoma differentiation by transcriptional repression of TERT. Mol Carcinog, 58(11), 1998–2007. doi:10.1002/mc.23091

Cancer Genome Atlas Research, N., Kandoth, C., Schultz, N., Cherniack, A. D., Akbani, R., Liu, Y., . . . Levine, D. A. (2013). Integrated genomic characterization of endometrial carcinoma. Nature, 497(7447), 67–73. doi:10.1038/nature12113

Chandler, R. L., Brennan, J., Schisler, J. C., Serber, D., Patterson, C., & Magnuson, T. (2013). ARID1a-DNA interactions are required for promoter occupancy by SWI/SNF. Molecular and Cellular Biology, 33(2), 265–280. doi:10.1128/MCB.01008-12

Chen, P., Zhao, J., Wang, Y., Wang, M., Long, H., Liang, D., . . . Li, G. (2013). H3.3 actively marks enhancers and primes gene transcription via opening higher-ordered chromatin. Genes Dev, 27(19), 2109–2124. doi:10.1101/gad.222174.113

Cheneby, J., Menetrier, Z., Mestdagh, M., Rosnet, T., Douida, A., Rhalloussi, W., . . . Ballester, A. (2020). ReMap 2020: a database of regulatory regions from an integrative analysis of Human and Arabidopsis DNA-binding sequencing experiments. Nucleic Acids Res, 48(D1), D180–D188. doi:10.1093/nar/gkz945

Clapier, C. R. (2021). Sophisticated Conversations between Chromatin and Chromatin Remodelers, and Dissonances in Cancer. Int J Mol Sci, 22(11). doi:10.3390/ijms22115578

Clapier, C. R., & Cairns, B. R. (2009). The biology of chromatin remodeling complexes. Annu Rev Biochem, 78, 273–304. doi:10.1146/annurev.biochem.77.062706.153223

Colino-Sanguino, Y., Cornett, E. M., Moulder, D., Smith, G. C., Hrit, J., Cordeiro-Spinetti, E., . . . Valdes-Mora, F. (2019). A Read/Write Mechanism Connects p300 Bromodomain Function to H2A.Z Acetylation. iScience, 21, 773–788. doi:10.1016/j.isci.2019.10.053

Cornett, E. M., Dickson, B. M., & Rothbart, S. B. (2017). Analysis of Histone Antibody Specificity with Peptide Microarrays. J Vis Exp(126). doi:10.3791/55912

Daley, T., & Smith, A. D. (2013). Predicting the molecular complexity of sequencing libraries. Nat Methods, 10(4), 325–327. doi:10.1038/nmeth.2375

Dallas, P. B., Pacchione, S., Wilsker, D., Bowrin, V., Kobayashi, R., & Moran, E. (2000). The human SWI-SNF complex protein p270 is an ARID family member with non-sequence-specific DNA binding activity. Mol Cell Biol, 20(9), 3137–3146. doi:10.1128/mcb.20.9.3137-3146.2000

Davis, S., & Meltzer, P. S. (2007). GEOquery: a bridge between the Gene Expression Omnibus (GEO) and BioConductor. Bioinformatics, 23(14), 1846–1847. doi:10.1093/bioinformatics/btm254

Deaton, A. M., Gomez-Rodriguez, M., Mieczkowski, J., Tolstorukov, M. Y., Kundu, S., Sadreyev, R. I., . . . Kingston, R. E. (2016). Enhancer regions show high histone H3.3 turnover that changes during differentiation. Elife, 5. doi:10.7554/eLife.15316

Dechassa, M. L., Sabri, A., Pondugula, S., Kassabov, S. R., Chatterjee, N., Kladde, M. P., & Bartholomew, B. (2010). SWI/SNF has intrinsic nucleosome disassembly activity that is dependent on adjacent nucleosomes. Mol Cell, 38(4), 590–602. doi:10.1016/j.molcel.2010.02.040

Dickson, B. M., Cornett, E. M., Ramjan, Z., & Rothbart, S. B. (2016). ArrayNinja: An Open Source Platform for Unified Planning and Analysis of Microarray Experiments. Methods Enzymol, 574, 53–77. doi:10.1016/bs.mie.2016.02.002

Dobin, A., Davis, C. A., Schlesinger, F., Drenkow, J., Zaleski, C., Jha, S., . . . Gingeras, T. R. (2013). STAR: ultrafast universal RNA-seq aligner. Bioinformatics, 29(1), 15–21. doi:10.1093/bioinformatics/bts635

Ernst, J., & Kellis, M. (2012). ChromHMM: automating chromatin-state discovery and characterization. Nature methods, 9(3), 215–216. doi:10.1038/nmeth.1906

Ewels, P., Magnusson, M., Lundin, S., & Kaller, M. (2016). MultiQC: summarize analysis results for multiple tools and samples in a single report. Bioinformatics, 32(19), 3047–3048. doi:10.1093/bioinformatics/btw354

Gehre, M., Bunina, D., Sidoli, S., Lubke, M. J., Diaz, N., Trovato, M., . . . Noh, K. M. (2020). Lysine 4 of histone H3.3 is required for embryonic stem cell differentiation, histone enrichment at regulatory regions and transcription accuracy. Nat Genet, 52(3), 273–282. doi:10.1038/s41588-020-0586-5

Ghandi, M., Huang, F. W., Jane-Valbuena, J., Kryukov, G. V., Lo, C. C., McDonald, E. R., 3rd, . . . Sellers, W. R. (2019). Next-generation characterization of the Cancer Cell Line Encyclopedia. Nature, 569(7757), 503–508. doi:10.1038/s41586-019-1186-3

Ghosh, K., Tang, M., Kumari, N., Nandy, A., Basu, S., Mall, D. P., . . . Biswas, D. (2018). Positive Regulation of Transcription by Human ZMYND8 through Its Association with P-TEFb Complex. Cell Reports, 24(8), 2141–2154 e2146. doi:10.1016/j.celrep.2018.07.064

Gong, F., Chiu, L. Y., Cox, B., Aymard, F., Clouaire, T., Leung, J. W., . . . Miller, K. M. (2015). Screen identifies bromodomain protein ZMYND8 in chromatin recognition of transcription-associated DNA damage that promotes homologous recombination. Genes Dev, 29(2), 197–211. doi:10.1101/gad.252189.114

Gong, F., Clouaire, T., Aguirrebengoa, M., Legube, G., & Miller, K. M. (2017). Histone demethylase KDM5A regulates the ZMYND8-NuRD chromatin remodeler to promote DNA repair. Journal of Cell Biology, 216(7), 1959–1974. doi:10.1083/jcb.201611135

Gorkin, D. U., Barozzi, I., Zhao, Y., Zhang, Y., Huang, H., Lee, A. Y., . . . Ren, B. (2020). An atlas of dynamic chromatin landscapes in mouse fetal development. Nature, 583(7818), 744–751. doi:10.1038/s41586-020-2093-3

Grandi, G., Toss, A., Cortesi, L., Botticelli, L., Volpe, A., & Cagnacci, A. (2015). The Association between Endometriomas and Ovarian Cancer: Preventive Effect of Inhibiting Ovulation and Menstruation during Reproductive Life. Biomed Res Int, 2015, 751571. doi:10.1155/2015/751571

Gu, Z., Eils, R., & Schlesner, M. (2016). Complex heatmaps reveal patterns and correlations in multidimensional genomic data. Bioinformatics, 32(18), 2847–2849. doi:10.1093/bioinformatics/btw313

Guo, R., Zheng, L., Park, J. W., Lv, R., Chen, H., Jiao, F., . . . Shi, Y. (2014). BS69/ZMYND11 reads and connects histone H3.3 lysine 36 trimethylation-decorated chromatin to regulated pre-mRNA processing. Molecular Cell, 56(2), 298–310. doi:10.1016/j.molcel.2014.08.022

Ha, M., Kraushaar, D. C., & Zhao, K. (2014). Genome-wide analysis of H3.3 dissociation reveals high nucleosome turnover at distal regulatory regions of embryonic stem cells. Epigenetics Chromatin, 7(1), 38. doi:10.1186/1756-8935-7-38

Hawkins, S. M., Creighton, C. J., Han, D. Y., Zariff, A., Anderson, M. L., Gunaratne, P. H., & Matzuk, M. M. (2011). Functional microRNA involved in endometriosis. Mol Endocrinol, 25(5), 821–832. doi:10.1210/me.2010-0371

Hazan, I., Monin, J., Bouwman, B. A. M., Crosetto, N., & Aqeilan, R. I. (2019). Activation of Oncogenic Super-Enhancers Is Coupled with DNA Repair by RAD51. Cell Rep, 29(3), 560–572 e564. doi:10.1016/j.celrep.2019.09.001

He, S., Wu, Z., Tian, Y., Yu, Z., Yu, J., Wang, X., . . . Xu, Y. (2020). Structure of nucleosome-bound human BAF complex. Science, 367(6480), 875–881. doi:10.1126/science.aaz9761

Heinz, S., Benner, C., Spann, N., Bertolino, E., Lin, Y. C., Laslo, P., . . . Glass, C. K. (2010). Simple combinations of lineage-determining transcription factors prime cis-regulatory elements required for macrophage and B cell identities. Mol Cell, 38(4), 576–589. doi:10.1016/j.molcel.2010.05.004

Ignatiadis, N., Klaus, B., Zaugg, J. B., & Huber, W. (2016). Data-driven hypothesis weighting increases detection power in genome-scale multiple testing. Nat. Methods, 13(7), 577–580. doi:10.1038/nmeth.3885

Jiao, F., Li, Z., He, C., Xu, W., Yang, G., Liu, T., . . . Guo, R. (2020). RACK7 recognizes H3.3G34R mutation to suppress expression of MHC class II complex components and their delivery pathway in pediatric glioblastoma. Sci Adv, 6(29), eaba2113. doi:10.1126/sciadv.aba2113

Jones, S., Wang, T. L., Shih Ie, M., Mao, T. L., Nakayama, K., Roden, R., . . . Papadopoulos, N. (2010). Frequent mutations of chromatin remodeling gene ARID1A in ovarian clear cell carcinoma. Science, 330(6001), 228–231. doi:10.1126/science.1196333

Kadoch, C., & Crabtree, G. R. (2015). Mammalian SWI/SNF chromatin remodeling complexes and cancer: Mechanistic insights gained from human genomics. Sci Adv, 1(5), e1500447. doi:10.1126/sciadv.1500447

Kadoch, C., Hargreaves, D. C., Hodges, C., Elias, L., Ho, L., Ranish, J., & Crabtree, G. R. (2013). Proteomic and bioinformatic analysis of mammalian SWI/SNF complexes identifies extensive roles in human malignancy. Nature Genetics, 45(6), 592–601. doi:10.1038/ng.2628

Kassabov, S. R., Zhang, B., Persinger, J., & Bartholomew, B. (2003). SWI/SNF unwraps, slides, and rewraps the nucleosome. Mol Cell, 11(2), 391–403. doi:10.1016/s1097-2765(03)00039-x

Kelso, T. W. R., Porter, D. K., Amaral, M. L., Shokhirev, M. N., Benner, C., & Hargreaves, D. C. (2017). Chromatin accessibility underlies synthetic lethality of SWI/SNF subunits in ARID1A-mutant cancers. Elife, 6. doi:10.7554/eLife.30506

Konev, A. Y., Tribus, M., Park, S. Y., Podhraski, V., Lim, C. Y., Emelyanov, A. V., . . . Fyodorov, D. V. (2007). CHD1 motor protein is required for deposition of histone variant H3.3 into chromatin in vivo. Science, 317(5841), 1087–1090. doi:10.1126/science.1145339

Kraushaar, D. C., Chen, Z., Tang, Q., Cui, K., Zhang, J., & Zhao, K. (2018). The gene repressor complex NuRD interacts with the histone variant H3.3 at promoters of active genes. Genome Res, 28(11), 1646–1655. doi:10.1101/gr.236224.118

Landt, S. G., Marinov, G. K., Kundaje, A., Kheradpour, P., Pauli, F., Batzoglou, S., . . . Snyder, M. (2012). ChIP-seq guidelines and practices of the ENCODE and modENCODE consortia. Genome Res, 22(9), 1813–1831. doi:10.1101/gr.136184.111

Langmead, B., & Salzberg, S. L. (2012). Fast gapped-read alignment with Bowtie 2. Nat Methods, 9(4), 357–359. doi:10.1038/nmeth.1923

Lawrence, M., Huber, W., Pages, H., Aboyoun, P., Carlson, M., Gentleman, R., . . . Carey, V. J. (2013). Software for computing and annotating genomic ranges. PLoS Comput Biol, 9(8), e1003118. doi:10.1371/journal.pcbi.1003118

Li, H., Handsaker, B., Wysoker, A., Fennell, T., Ruan, J., Homer, N., . . . Genome Project Data Processing, S. (2009). The Sequence Alignment/Map format and SAMtools. Bioinformatics, 25(16), 2078–2079. doi:10.1093/bioinformatics/btp352

Li, N., Li, Y., Lv, J., Zheng, X., Wen, H., Shen, H., . . . Lee, M. G. (2016). ZMYND8 Reads the Dual Histone Mark H3K4me1-H3K14ac to Antagonize the Expression of Metastasis-Linked Genes. Molecular Cell, 63(3), 470–484. doi:10.1016/j.molcel.2016.06.035

Local, A., Huang, H., Albuquerque, C. P., Singh, N., Lee, A. Y., Wang, W., . . . Ren, B. (2018). Identification of H3K4me1-associated proteins at mammalian enhancers. Nat Genet, 50(1), 73–82. doi:10.1038/s41588-017-0015-6

Love, M. I., Anders, S., Kim, V., & Huber, W. (2015). RNA-Seq workflow: gene-level exploratory analysis and differential expression. F1000Res, 4, 1070. doi:10.12688/f1000research.7035.1

Love, M. I., Huber, W., & Anders, S. (2014). Moderated estimation of fold change and dispersion for RNA-seq data with DESeq2. Genome Biology, 15(12), 550. doi:10.1186/s13059-014-0550-8

Lun, A. T., & Smyth, G. K. (2016). csaw: a Bioconductor package for differential binding analysis of ChIP-seq data using sliding windows. Nucleic Acids Res, 44(5), e45. doi:10.1093/nar/gkv1191

Marques, J. G., Gryder, B. E., Pavlovic, B., Chung, Y., Ngo, Q. A., Frommelt, F., . . . Schafer, B. W. (2020). NuRD subunit CHD4 regulates super-enhancer accessibility in rhabdomyosarcoma and represents a general tumor dependency. Elife, 9. doi:10.7554/eLife.54993

Martin, M. (2011). Cutadapt removes adapter sequences from high-throughput sequencing reads. EMBnet. J., 17, 10–12.

Martire, S., Gogate, A. A., Whitmill, A., Tafessu, A., Nguyen, J., Teng, Y. C., . . . Banaszynski, L. A. (2019). Phosphorylation of histone H3.3 at serine 31 promotes p300 activity and enhancer acetylation. Nat Genet, 51(6), 941–946. doi:10.1038/s41588-019-0428-5

Mashtalir, N., D’Avino, A. R., Michel, B. C., Luo, J., Pan, J., Otto, J. E., . . . Kadoch, C. (2018). Modular Organization and Assembly of SWI/SNF Family Chromatin Remodeling Complexes. Cell, 175(5), 1272–1288 e1220. doi:10.1016/j.cell.2018.09.032

Mittal, P., & Roberts, C. W. M. (2020). The SWI/SNF complex in cancer - biology, biomarkers and therapy. Nature Reviews: Clinical Oncology, 17(7), 435–448. doi:10.1038/s41571-020-0357-3

Morris, S. A., Baek, S., Sung, M. H., John, S., Wiench, M., Johnson, T. A., . . . Hager, G. L. (2014). Overlapping chromatin-remodeling systems collaborate genome wide at dynamic chromatin transitions. Nat Struct Mol Biol, 21(1), 73–81. doi:10.1038/nsmb.2718

Nacev, B. A., Feng, L., Bagert, J. D., Lemiesz, A. E., Gao, J., Soshnev, A. A., . . . Allis, C. D. (2019). The expanding landscape of ‘oncohistone’ mutations in human cancers. Nature, 567(7749), 473–478. doi:10.1038/s41586-019-1038-1

Ou, J., Liu, H., Yu, J., Kelliher, M. A., Castilla, L. H., Lawson, N. D., & Zhu, L. J. (2018). ATACseqQC: a Bioconductor package for post-alignment quality assessment of ATAC-seq data. BMC Genomics, 19(1), 169. doi:10.1186/s12864-018-4559-3

Park, J. H., Park, E. J., Lee, H. S., Kim, S. J., Hur, S. K., Imbalzano, A. N., & Kwon, J. (2006). Mammalian SWI/SNF complexes facilitate DNA double-strand break repair by promoting gamma-H2AX induction. Embo J, 25(17), 3986–3997. doi:10.1038/sj.emboj.7601291

Pillidge, Z., & Bray, S. J. (2019). SWI/SNF chromatin remodeling controls Notch-responsive enhancer accessibility. EMBO Rep, 20(5). doi:10.15252/embr.201846944

Pradhan, S. K., Su, T., Yen, L., Jacquet, K., Huang, C., Cote, J., . . . Carey, M. F. (2016). EP400 Deposits H3.3 into Promoters and Enhancers during Gene Activation. Mol Cell, 61(1), 27–38. doi:10.1016/j.molcel.2015.10.039

Quinlan, A. R., & Hall, I. M. (2010). BEDTools: a flexible suite of utilities for comparing genomic features. Bioinformatics, 26(6), 841–842. doi:10.1093/bioinformatics/btq033

R Core Team. (2018). R: A language and environment for statistical computing. Retrieved from https://www.R-project.org/

Rafati, H., Parra, M., Hakre, S., Moshkin, Y., Verdin, E., & Mahmoudi, T. (2011). Repressive LTR nucleosome positioning by the BAF complex is required for HIV latency. PLoS Biol, 9(11), e1001206. doi:10.1371/journal.pbio.1001206

Reske, J. J., Wilson, M. R., Holladay, J., Wegener, M., Adams, M., & Chandler, R. L. (2020). SWI/SNF inactivation in the endometrial epithelium leads to loss of epithelial integrity. Human Molecular Genetics, 29(20), 3412–3430. doi:10.1093/hmg/ddaa227

Ritchie, M. E., Phipson, B., Wu, D., Hu, Y., Law, C. W., Shi, W., & Smyth, G. K. (2015). limma powers differential expression analyses for RNA-sequencing and microarray studies. Nucleic Acids Res, 43(7), e47. doi:10.1093/nar/gkv007

Robinson, J. T., Thorvaldsdottir, H., Winckler, W., Guttman, M., Lander, E. S., Getz, G., & Mesirov, J. P. (2011). Integrative genomics viewer. Nat Biotechnol, 29(1), 24–26. doi:10.1038/nbt.1754

Robinson, M. D., McCarthy, D. J., & Smyth, G. K. (2010). edgeR: a Bioconductor package for differential expression analysis of digital gene expression data. Bioinformatics, 26(1), 139–140. doi:10.1093/bioinformatics/btp616

Rothbart, S. B., Krajewski, K., Strahl, B. D., & Fuchs, S. M. (2012). Peptide microarrays to interrogate the “histone code”. Methods Enzymol, 512, 107–135. doi:10.1016/B978-0-12-391940-3.00006-8

Samartzis, E. P., Samartzis, N., Noske, A., Fedier, A., Caduff, R., Dedes, K. J., . . . Imesch, P. (2012). Loss of ARID1A/BAF250a-expression in endometriosis: a biomarker for risk of carcinogenic transformation? Modern Pathology, 25(6), 885–892. doi:10.1038/modpathol.2011.217

Savitsky, P., Krojer, T., Fujisawa, T., Lambert, J. P., Picaud, S., Wang, C. Y., . . . Filippakopoulos, P. (2016). Multivalent Histone and DNA Engagement by a PHD/BRD/PWWP Triple Reader Cassette Recruits ZMYND8 to K14ac-Rich Chromatin. Cell Rep, 17(10), 2724–2737. doi:10.1016/j.celrep.2016.11.014

Schneider, V. A., Graves-Lindsay, T., Howe, K., Bouk, N., Chen, H. C., Kitts, P. A., . . . Church, D. M. (2017). Evaluation of GRCh38 and de novo haploid genome assemblies demonstrates the enduring quality of the reference assembly. Genome Res, 27(5), 849–864. doi:10.1101/gr.213611.116

Schwabish, M. A., & Struhl, K. (2007). The Swi/Snf complex is important for histone eviction during transcriptional activation and RNA polymerase II elongation in vivo. Mol Cell Biol, 27(20), 6987–6995. doi:10.1128/MCB.00717-07

Serresi, M., Kertalli, S., Li, L., Schmitt, M. J., Dramaretska, Y., Wierikx, J., . . . Gargiulo, G. (2021). Functional antagonism of chromatin modulators regulates epithelial-mesenchymal transition. Sci Adv, 7(9). doi:10.1126/sciadv.abd7974

Shen, H., Xu, W., Guo, R., Rong, B., Gu, L., Wang, Z., . . . Lan, F. (2016). Suppression of Enhancer Overactivation by a RACK7-Histone Demethylase Complex. Cell, 165(2), 331–342. doi:10.1016/j.cell.2016.02.064

Shi, L., Wen, H., & Shi, X. (2017). The Histone Variant H3.3 in Transcriptional Regulation and Human Disease. J Mol Biol, 429(13), 1934–1945. doi:10.1016/j.jmb.2016.11.019

Sikora, J., Wroblewska-Czech, A., Smycz-Kubanska, M., Mielczarek-Palacz, A., Cygal, A., Witek, A., & Kondera-Anasz, Z. (2018). The role of complement components C1q, MBL and C1 inhibitor in pathogenesis of endometriosis. Arch Gynecol Obstet, 297(6), 1495–1501. doi:10.1007/s00404-018-4754-0

Smedley, D., Haider, S., Ballester, B., Holland, R., London, D., Thorisson, G., & Kasprzyk, A. (2009). BioMart--biological queries made easy. BMC Genomics, 10, 22. doi:10.1186/1471-2164-10-22

Smeenk, G., Wiegant, W. W., Vrolijk, H., Solari, A. P., Pastink, A., & van Attikum, H. (2010). The NuRD chromatin-remodeling complex regulates signaling and repair of DNA damage. J Cell Biol, 190(5), 741–749. doi:10.1083/jcb.201001048

Spruijt, C. G., Luijsterburg, M. S., Menafra, R., Lindeboom, R. G., Jansen, P. W., Edupuganti, R. R., . . . Vermeulen, M. (2016). ZMYND8 Co-localizes with NuRD on Target Genes and Regulates Poly(ADP-Ribose)-Dependent Recruitment of GATAD2A/NuRD to Sites of DNA Damage. Cell Rep, 17(3), 783–798. doi:10.1016/j.celrep.2016.09.037

Szenker, E., Ray-Gallet, D., & Almouzni, G. (2011). The double face of the histone variant H3.3. Cell Res, 21(3), 421–434. doi:10.1038/cr.2011.14

Tong, J. K., Hassig, C. A., Schnitzler, G. R., Kingston, R. E., & Schreiber, S. L. (1998). Chromatin deacetylation by an ATP-dependent nucleosome remodelling complex. Nature, 395(6705), 917–921. doi:10.1038/27699

Trizzino, M., Barbieri, E., Petracovici, A., Wu, S., Welsh, S. A., Owens, T. A., . . . Gardini, A. (2018). The Tumor Suppressor ARID1A Controls Global Transcription via Pausing of RNA Polymerase II. Cell Reports, 23(13), 3933–3945. doi:10.1016/j.celrep.2018.05.097

Van Rechem, C., Boulay, G., & Leprince, D. (2009). HIC1 interacts with a specific subunit of SWI/SNF complexes, ARID1A/BAF250A. Biochem Biophys Res Commun, 385(4), 586–590. doi:10.1016/j.bbrc.2009.05.115

Wang, W., Cote, J., Xue, Y., Zhou, S., Khavari, P. A., Biggar, S. R., . . . Crabtree, G. R. (1996). Purification and biochemical heterogeneity of the mammalian SWI-SNF complex. EMBO J, 15(19), 5370–5382. Retrieved from https://www.ncbi.nlm.nih.gov/pubmed/8895581 https://www.ncbi.nlm.nih.gov/pmc/articles/PMC452280/pdf/emboj00019-0250.pdf

Wang, W., Vilella, F., Alama, P., Moreno, I., Mignardi, M., Isakova, A., . . . Quake, S. R. (2020). Single-cell transcriptomic atlas of the human endometrium during the menstrual cycle. Nat Med, 26(10), 1644–1653. doi:10.1038/s41591-020-1040-z

Wen, H., Li, Y., Xi, Y., Jiang, S., Stratton, S., Peng, D., . . . Shi, X. (2014). ZMYND11 links histone H3.3K36me3 to transcription elongation and tumour suppression. Nature, 508(7495), 263–268. doi:10.1038/nature13045

Whitehouse, I., Flaus, A., Cairns, B. R., White, M. F., Workman, J. L., & Owen-Hughes, T. (1999). Nucleosome mobilization catalysed by the yeast SWI/SNF complex. Nature, 400(6746), 784–787. doi:10.1038/23506

Whyte, W. A., Orlando, D. A., Hnisz, D., Abraham, B. J., Lin, C. Y., Kagey, M. H., . . . Young, R. A. (2013). Master transcription factors and mediator establish super-enhancers at key cell identity genes. Cell, 153(2), 307–319. doi:10.1016/j.cell.2013.03.035

Wickham, H. (2016). ggplot2: Elegant Graphics for Data Analysis: Springer-Verlag New York.

Wiegand, K. C., Shah, S. P., Al-Agha, O. M., Zhao, Y., Tse, K., Zeng, T., . . . Huntsman, D. G. (2010). ARID1A mutations in endometriosis-associated ovarian carcinomas. N Engl J Med, 363(16), 1532–1543. doi:10.1056/NEJMoa1008433

Wilson, M. R., Reske, J. J., Holladay, J., Neupane, S., Ngo, J., Cuthrell, N., . . . Chandler, R. L. (2020). ARID1A Mutations Promote P300-Dependent Endometrial Invasion through Super-Enhancer Hyperacetylation. Cell Reports, 33(6), 108366. doi:10.1016/j.celrep.2020.108366

Wilson, M. R., Reske, J. J., Holladay, J., Wilber, G. E., Rhodes, M., Koeman, J., . . . Chandler, R. L. (2019). ARID1A and PI3-kinase pathway mutations in the endometrium drive epithelial transdifferentiation and collective invasion. Nat Commun, 10(1), 3554. doi:10.1038/s41467-019-11403-6

Wu, R. C., Wang, T. L., & Shih Ie, M. (2014). The emerging roles of ARID1A in tumor suppression. Cancer Biology & Therapy, 15(6), 655–664. doi:10.4161/cbt.28411

Xue, Y., Wong, J., Moreno, G. T., Young, M. K., Cote, J., & Wang, W. (1998). NURD, a novel complex with both ATP-dependent chromatin-remodeling and histone deacetylase activities. Mol Cell, 2(6), 851–861. doi:10.1016/s1097-2765(00)80299-3

Zhang, C., Wang, X., Anaya, Y., Parodi, L., Cheng, L., Anderson, M. L., & Hawkins, S. M. (2018). Distinct molecular pathways in ovarian endometrioid adenocarcinoma with concurrent endometriosis. Int J Cancer, 143(10), 2505–2515. doi:10.1002/ijc.31768

Zhang, H., Gan, H., Wang, Z., Lee, J. H., Zhou, H., Ordog, T., . . . Zhang, Z. (2017). RPA Interacts with HIRA and Regulates H3.3 Deposition at Gene Regulatory Elements in Mammalian Cells. Mol Cell, 65(2), 272–284. doi:10.1016/j.molcel.2016.11.030

Zhang, Y., Liu, T., Meyer, C. A., Eeckhoute, J., Johnson, D. S., Bernstein, B. E., . . . Liu, X. S. (2008). Model-based analysis of ChIP-Seq (MACS). Genome Biol, 9(9), R137. doi:10.1186/gb-2008-9-9-r137

Zhou, Y., & Grummt, I. (2005). The PHD finger/bromodomain of NoRC interacts with acetylated histone H4K16 and is sufficient for rDNA silencing. Curr Biol, 15(15), 1434–1438. doi:10.1016/j.cub.2005.06.057

Zhu, A., Ibrahim, J. G., & Love, M. I. (2019). Heavy-tailed prior distributions for sequence count data: removing the noise and preserving large differences. Bioinformatics, 35(12), 2084–2092. doi:10.1093/bioinformatics/bty895

Zondervan, K. T., Becker, C. M., & Missmer, S. A. (2020). Endometriosis. N Engl J Med, 382(13), 1244–1256. doi:10.1056/NEJMra1810764

